# Functional prediction of proteins from the human gut archaeome

**DOI:** 10.1101/2023.02.01.526569

**Authors:** Polina V. Novikova, Susheel Bhanu Busi, Alexander J. Probst, Patrick May, Paul Wilmes

## Abstract

The human gastrointestinal tract contains diverse microbial communities, including archaea. Among them, *Methanobrevibacter smithii* represents a highly active and clinically relevant methanogenic archaeon, being involved in gastrointestinal disorders, such as IBD and obesity. Herein, we present an integrated approach using sequence and structure information to improve the annotation of *M. smithii* proteins using advanced protein structure prediction and annotation tools, such as AlphaFold2, trRosetta, ProFunc, and DeepFri. Of an initial set of 873 481 archaeal proteins, we found 707 754 proteins exclusively present in the human gut. Having analysed archaeal proteins together with 87 282 994 bacterial proteins, we identified unique archaeal proteins and archaeal-bacterial homologs. We then predicted and characterized functional domains and structures of 73 unique and homologous archaeal protein clusters linked the human gut and *M. smithii*. We refined annotations based on the predicted structures, extending existing sequence similarity-based annotations. We identified gut-specific archaeal proteins that may be involved in defense mechanisms, virulence, adhesion, and the degradation of toxic substances. Interestingly, we identified potential glycosyltransferases that could be associated with N-linked and O-glycosylation. Additionally, we found preliminary evidence for interdomain horizontal gene transfer between *Clostridia* species and *M. smithii*, which includes *sporulation stage V proteins AE* and AD. Our study broadens the understanding of archaeal biology, particularly *M. smithii,* and highlights the importance of considering both sequence and structure for the prediction of protein function.

## Introduction

In 1977, Woese and Fox, and colleagues discovered the kingdom of Archaebacteria, later renamed Archaea, revealing a new branch in the tree of life [1]–[4]. The discovery of the Asgard superphylum and its close relationship with the eukaryotic branch supports the notion of an archaeal origin for eukaryotes, yet ongoing debates continue regarding whether the archaeal ancestor of eukaryotes belongs within the Asgard superphylum or represents a sister group to all other archaea [5], [6]. Historically, archaea were associated with extreme environments but have since been recognized for their general importance and prevalence [7], [8]. Their ability to thrive in extreme environments and to resist chemicals is attributed, in part, to their unique cell envelope structures. In nature, archaea perform distinctive biogeochemical functions, such as methanogenesis, anaerobic methane oxidation, and ammonia oxidation [9], [10]. By employing diverse ecological strategies for energy production, archaea can inhabit a wide variety of environments [11]. Archaea are also host-associated, such as on plants, in human and animal gastrointestinal tracts [12], [13], on human skin [14], [15], in respiratory airways [16] and in the oral cavity [17]. Based on recent estimates, archaea comprise up to 10% of the human gut microbiota [18].

*Methanobrevibacter smithii*, a ubiquitous and highly active methanogen in the human gut microbiome, has remarkable clinical relevance and is relatively well annotated [19]. It plays an important role in the degradation of complex carbohydrates, leading to the production of methane, which has significant physiological effects on human physiology. Imbalances in the population of *M. smithii* have been implicated as factors contributing to gastrointestinal disorders such as inflammatory bowel disease (IBD) [20], [21] and obesity [22]–[24]. Given the prevalence of *M. smithii* in the gut, further research aimed at *M. smithii* is key to understanding their role in disease. Archaeal proteins, including those of *M. smithii*, play a crucial role in adapting to diverse environments and showcase their unique biology. The knowledge about diverse archaea, including novel species, in the human gut microbiome has expanded, underscoring their significance [25]. Some host-associated taxa, like *Methanomassilicoccales*, have potential beneficial effects on human health [26], while others like *Methanosphaera stadtmanae* have been linked to pro-inflammatory immune processes [27]. Given the current interest in the role of archaea in human health and disease, understanding the archaeal proteome is crucial for understanding the functional potential of archaea.

Studying archaeal proteins presents challenges both in experimental and computational aspects. Previous research by Lyu *et al.* has highlighted the potential for biotechnological applications in various archaeal genera [28]. However, genetic toolboxes for targeted genomic modifications are currently limited to mesophilic *Methanococcus* and *Methanosarcina* genera [29]. While alternative methods like mass spectrometry-based searches exist, difficulties arise from inaccurate predictions of protein coding sequences due to limited knowledge of ribosomal binding sites and promoter consensus sequences [30]. Another unresolved challenge lies in the isolation and cultivation of archaea under laboratory conditions, although recent progress has been made in this area [31], [32]. To overcome these challenges, metagenomic sequencing has emerged as a promising approach to study archaea and their ecological relationships. Metagenomics has enhanced our understanding of the archaeal branches within the tree of life [31]–[33] whereby assembled sequences allow prediction of protein coding sequences (CDS) and their functional characterization *in silico*. However, metagenome-assembled genomes (MAGs) face challenges in functional assignment due to incomplete sequences and difficulties in predicting and annotating open-reading frames (ORFs) [34], [35]. Sequence-based protein function annotation, commonly used but limited in cases of distant protein homologies, proves to be not particularly effective [36]. Moreover, the databases containing information about archaeal proteins and functions are not consistently updated, creating a twofold challenge in the sequence-based annotation of archaeal proteins. On one hand, Makarova *et al*. [37] report that *archaeal ribosomal proteins L45* and L47, experimentally identified in 2011 [38] and pre-rRNA processing and ribosome biogenesis proteins of the *NOL1/NOP2/fmu* family characterized in 1998 [39] were not added to annotation pipelines by 2019 and were labelled as ‘hypothetical’. On the other hand, sequence similarity-based approaches fail to capture relationships between highly divergent proteins when aligned with a known database protein [40]–[42]. Archaea, the least characterized domain of life, suffer from spurious and incorrect protein annotations due to insufficient experimental data and outdated databases [43]. Furthermore, the study by Makarova *et al.* indicates that a substantial proportion of genes within archaeal genomes, estimated to be between 30% and 80%, have not been thoroughly characterized, leading to their classification as archaeal “dark matter” [37]. Poorly annotated proteins limit our study of microbial functionality and their roles in specific processes. However, protein structure prediction represents an alternative strategy to address the gap in sequence-function annotation [44]. It complements sequence-based approaches, particularly when annotations are limited or conflicting across databases, by utilizing the conservation of tertiary structure to infer functional roles [45], [46]. Advanced computational techniques, such as AlphaFold2 [47] and trRosetta [48], offer accurate predictions of three-dimensional structures, thus providing valuable functional insights.

In this study, we introduce an integrated *in silico* approach that aims to refine the functional characterization of proteins, thus enhancing the accuracy of protein annotations in the archaeon *M. smithii*. Having compared archaeal gut-specific proteins to bacterial gut proteins, we found 73 unique and homologous archaeal protein clusters. Our approach incorporates advanced protein structure prediction and annotation tools, such as AlphaFold2, trRosetta, ProFunc, and DeepFri, into a comprehensive workflow. We predict and characterize the functional domains and structures of 73 gut-specific archaeal protein clusters. The predicted functions are linked to the adaptation to changing environments, survival, and nutritional capabilities of *M. smithii* within the human gut microbiome. We additionally identified sporulation-related archaeal proteins, presumably horizontally transferred to archaea from *Clostridium* species.

## Materials and Methods

### Selection of gut-specific archaeal proteins

To select specific proteins of gut-associated archaea, we utilized archaeal MAGs obtained from the Genomes from Earth’s Microbiomes (GEM) catalog [49] and the Unified Human Gastrointestinal Genome (UHGG) collection [50], along with bacterial MAGs from the UHGG collection (accessed in November 2020). Genomes were extracted based on available metadata and filtered by taxonomy to specifically target archaea.

Gene prediction was performed using Prodigal (V2.6.3: February, 2016) [51] on the archaeal and bacterial MAGs from the UHGG collection, while CDSs from the GEM catalog were downloaded from the provided source (https://portal.nersc.gov/GEM). Archaeal and bacterial proteins were further separately clustered using MMseqs2 (MM2) (v12.113e3-2) [52], [53] (Fig. 1) with the following parameters: --cov-mode 0 --min-seq-id 0.9 -c 0.9.

**Figure 1.**
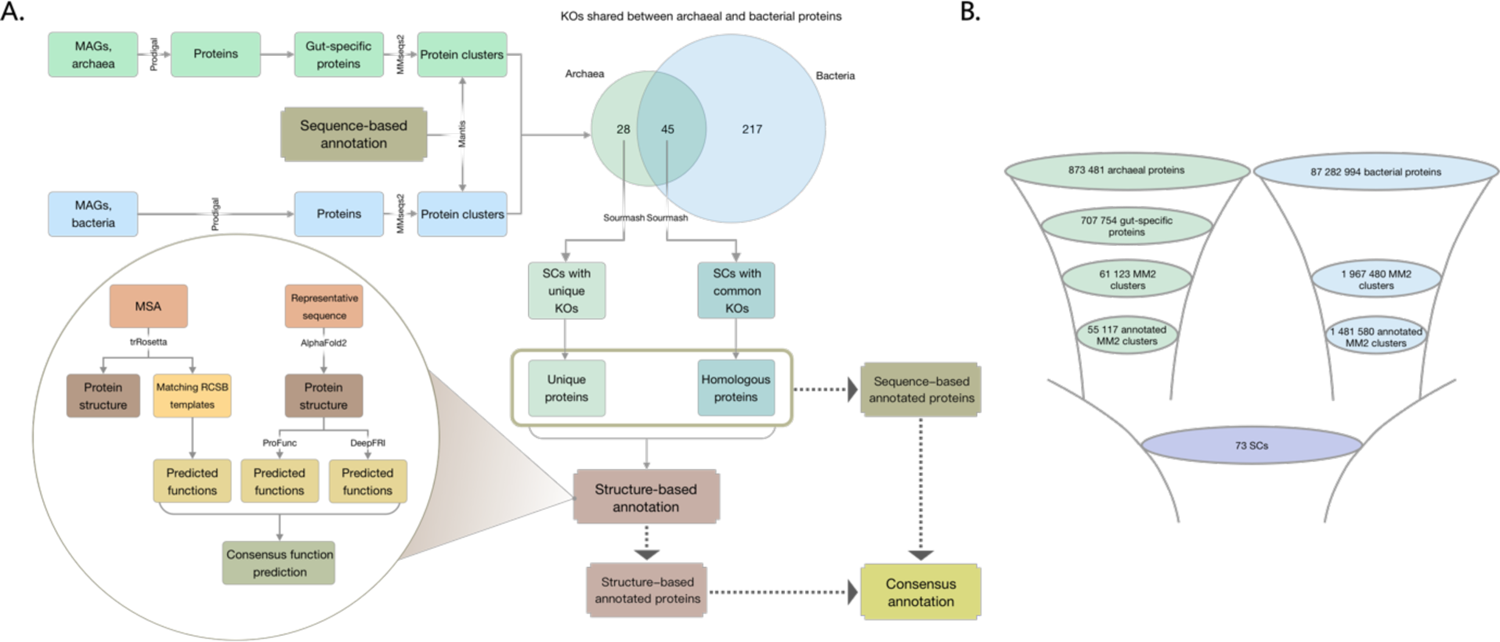
**A**, Flowchart demonstrating major steps of the analysis. The Venn diagram demonstrates the number of shared KOs assigned to archaeal and bacterial smash clusters. **B**, Funnels illustrating the protein count at each stage of protein selection. MM2 – Mmseqs2 clusters, SCs – smash clusters.

To identify unique functionalities of gut-associated archaea, we selected proteins that are specific to the human gut and encoded by gut-associated archaea. MAGs were selected based on available metadata indicating their sampling location. First, we included protein clusters containing at least one protein from a MAG sampled in the human gut. We then excluded protein clusters that had proteins from MAGs sampled in other environments, as these were outside the scope of the study. The final selection included protein clusters where all proteins were encoded by MAGs sampled exclusively from the human gut.

From the selected gut-specific protein clusters, only those with complete KEGG annotations were included. Fully annotated MM2 clusters were additionally clustered with Sourmash (v4.0.0) into sourmash clusters [54], [55]. Archaeal protein clusters were categorized into two groups: those sharing KEGG Orthology identifiers (KOs) with bacterial proteins (prefix *h*) and those with unique KOs (prefix *u*) (Fig. 1).

### Protein function annotation

Archaeal and bacterial proteins were annotated with KEGG orthologs (KOs) using Mantis (release 1.5.4) [56] (Fig. 1). AlphaFold2 (AF) [47], [57] and TrRosetta (TR) [48] were used as structure-prediction tools. For each tool, the predicted protein structure was then annotated separately. The TR-based model was annotated using templates with the highest *identity* and *coverage* features. TR used a template for prediction if it met the criteria of confidence > 0.6, e-value < 0.001, and coverage > 0.3. The protein model generated by AF was submitted to the ProFunc (PF) [58] web server for structure-based annotation. *Sequence search vs existing PDB entries* and 3D functional template searches sections from the ProFunc report were used for structure-based protein annotation. Structure matches were selected according to the reported highest possible likelihood of being correct as follows: *certain matches (E-value <10-6)*, *probable matches (10-6 < E-value <0.01)*, *possible matches (0.01 < E-value < 0.1)*, and *long shots (0.1 < E-value < 10.0)*. Only certain matches were used for the functional assignment. DeepFri [59] was used as an auxiliary tool, providing broad and general descriptions to verify or refute suggestions from AF and/or TR. DeepFri predictions with a certainty score > 0.7 were considered. Our combined approach integrates multiple methods to enhance the resolution of functional annotation, particularly for challenges faced by traditional sequence-based methods.

When TR- and AF-based annotations provided consistent results, the consensus was used as the final annotation of the protein function. However, when the reports gave different results, we prioritized the result with highest confidence. For instance, when the confidence of the model predicted by TR was *very high* and template matches were provided, and AF-based ProFunc reported a match with a lower confidence (anything but *certain match*), the template hit by TR was used as the primary source for the annotation. The relationship between PF likelihood and TR Template Modeling scores (TM-scores) generated in our analysis is shown in Table 1. Similarly, any protein with a TR template match was considered as more reliable than an annotation with the *long shot* likelihood. In cases where there were no 3D functional hits, TR annotation was given priority. In cases when PF and TR provided annotations with the same level of significance/likelihood, the protein structure with highest *coverage* and *identity* was chosen. In this case, we define coverage as *coverage* feature in TR and the 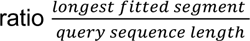 as in PF, and for identity we take *identity* as in

### TR and percentage sequence identity as in PF

The appropriateness of an annotation was determined based on the extent to which the assigned function of a protein was found to be directly relevant to archaea and supported by relevant literature. Any other annotations were classified as incorrect. Following this initial step, *sensitivity* was calculated as 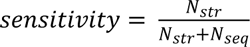, *specificity* as 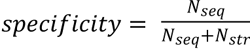, *positive likelihood ratio (PLR)* as 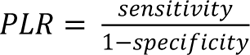, negative likelihood ratio (NLR) as 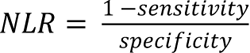 where N_seq_ and N_str_ are the numbers of correct sequence- and structure-based annotations, respectively.

### Protein relative occurrence calculation

Relative occurrence or frequency of protein functions in the groups of unique and homologous proteins was calculated. The measure was calculated as the ratio of the number of proteins with a specific KO to the total number of proteins of bacterial or archaeal proteins. For example, the relative occurrence of unique archaeal proteins annotated as K20411 (sourmash cluster 1) is: 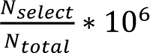, where N_selet_ is the amount of proteins annotated with K20411 and N_total_ is the total number of archaeal proteins. The reason for using a constant factor of 10^1^ in the equation is to scale the values and generate numbers better suited for graphical representation.

### Gene expression analysis

To assess the expression of archaeal proteins in the context of human health and disease, gene expression was verified using an in-house dataset, by mapping metatranscriptomic reads of faecal samples of healthy individuals and patients with type 1 diabetes mellitus (T1DM) [60] to nucleotide sequences of genes of interest using bwa mem [61]. Mapping files were processed with SAMtools (v1.6) [62]. Mosdepth (v0.3.3) [63] was used to calculate mean read coverage per gene of interest.

### Horizontal gene transfer analysis

To assess the stability of gene structures in *M. smithii* genomes, we conducted a horizontal gene transfer (HGT) analysis using metaCHIP (v1.10.12) [64] on all *M. smithii* MAGs available in the included datasets. One *Methanobrevibacter_A oralis* MAG derived from UHGG were also included for the comparison of the number of HGT events.

### Gene synteny analysis

pyGenomeViz (v0.3.2) [65] was used to build gene synteny for all archaeal genes of interest. Gene coordinates predicted with Prodigal were used as an input. An interval of 10kb up- and downstream of the gene of interest was selected from the protein predictions. KEGG KOs were allocated based on the sequence-based annotations generated using Mantis [56]. The *M. smithii* type strain DSM 861 was used to assess the presence of genes from flanking regions of specific genes in an archaeal culture. In our study, we exclusively focused on *M. smithii*, as our analysis revealed that all the gut-specific proteins encoded by gut-associated archaea were encoded by *M. smithii*.

### Phylogenetic analysis

In order to build phylogenetic trees for selective sourmash clusters, additional similar sequences were added from Uniprot [66] using BLAST (v2.0.15.153) [67] with default parameters on the consensus sequences representing sourmash clusters of interest, namely *h9* and *h20*. Furthermore, Uniprot sequences and sourmash cluster sequences were used to build trees. Multiple sequence alignments were built using MAFFT (v7) [68] and trimmed with BMGE (v1.12) [69] using BLOSUM95 similarity matrix and the default cut-off 0.5. Maximum likelihood phylogenetic trees were built with IQ-TREE (v1.6.12) [70] and visualized using the R library *ggtree* (v3.6.2) [71].

## Results & Discussion

Our study aimed to analyze the gut-specific proteins encoded by *M. smithii* in the human gastrointestinal tract. As we focused on identifying archaeal unique proteins and archaea-bacterial homologs, we analysed gut-specific archaeal and gut bacterial proteins together. Having compared the two subsets based on their sequence-based annotation, we categorized archaeal gut-specific proteins into two groups: unique and homologous proteins. To annotate them, we used KEGG KOs due to their consistent functional annotations across organisms and widespread usage. For structure-based functional assignment, we utilized a combination of structure prediction and annotation tools (Fig. 1), leveraging the higher prediction accuracy of AlphaFold2 and the rapid and accurate *de novo* predictions obtained via TrRosetta. Utilizing representative sequences of unique and homologous proteins, AlphaFold2 produced protein structures, and subsequent functional annotations were accomplished by integrating ProFunc and DeepFRI. trRosetta was employed to predict structures of unique and homologous proteins showing detectable homologous matches in the Protein Data Bank, which were subsequently used for further structure annotation.

### Enhancing annotations of proteins encoded by *Methanobrevibacter smithii*

To explore the uncharted functional space of *M. smithii*, we first selected gut-specific proteins of gut-associated archaea. We collected the encoded proteins of a total of 1 190 archaeal and 285 835 bacterial MAGs, resulting in 873 481 archaeal proteins and 87 282 994 bacterial proteins (Fig. 1). We focused on proteins associated with archaea of the human gut microbiome, which represented 37% (707 754 proteins) of all predicted archaeal proteins. These proteins were grouped into 61 123 MMseqs2 clusters for archaea (≥2 proteins per cluster) and 1 967 480 MM2 clusters for bacteria (≥10 proteins per cluster). By retaining fully annotated protein clusters, we obtained 55 117 archaeal MM2 clusters and 1 481 580 bacterial MM2 clusters. Using our proposed functional prediction strategy (Fig. 1A), we analyzed the gut-associated archaeal proteins alongside bacterial proteins, resulting in 45 homologous sourmash clusters, *i.e.*, shared between archaea and bacteria, and 28 unique sourmash clusters, *i.e.*, composed exclusively of archaeal proteins. The bacterial data served as a reference to distinguish unique proteins encoded and transcribed by archaea, as well as archaeal proteins with homologs to bacterial ORFs. A summary of the annotations is provided in the Supplementary Materials (Supp. Tab. 1-2).

All archaeal proteins from the abovementioned sourmash clusters were classified as *M. smithii*. We thus sought to extend our knowledge of *M. smithii* by exploring functions that could have implications for human health and disease. The investigation of the relative abundance of identified proteins and their associated processes revealed distinct types of functions in unique and homologous protein clusters (Fig. 2). The most frequently identified functions in the unique sourmash clusters were related to adaptation to changing environments and protection mechanisms, *e.g.,* defense against foreign DNA and oxidative stress, while processes such as RNA and DNA regulation, energy metabolism, and cell wall integrity and maintenance were less represented (Supp. Tab. 3). Homologous sourmash clusters showed frequent functions related to adaptation, various protection mechanisms, energy metabolism, and cell structural integrity (Supp. Tab. 4). Analysis of fecal metatranscriptomic data confirmed the transcription of the majority of encoded genes, with some unique and homologous genes exhibiting higher expression levels (Fig. 2). Two unique and 19 homologous sourmash clusters with relatively high expression levels were identified, including genes associated with adaptation to changing environments, defense against foreign DNA and oxidative stress, DNA/RNA regulation, and energy metabolism, while the rest were unannotated (Fig. 2).

**Figure 2.**
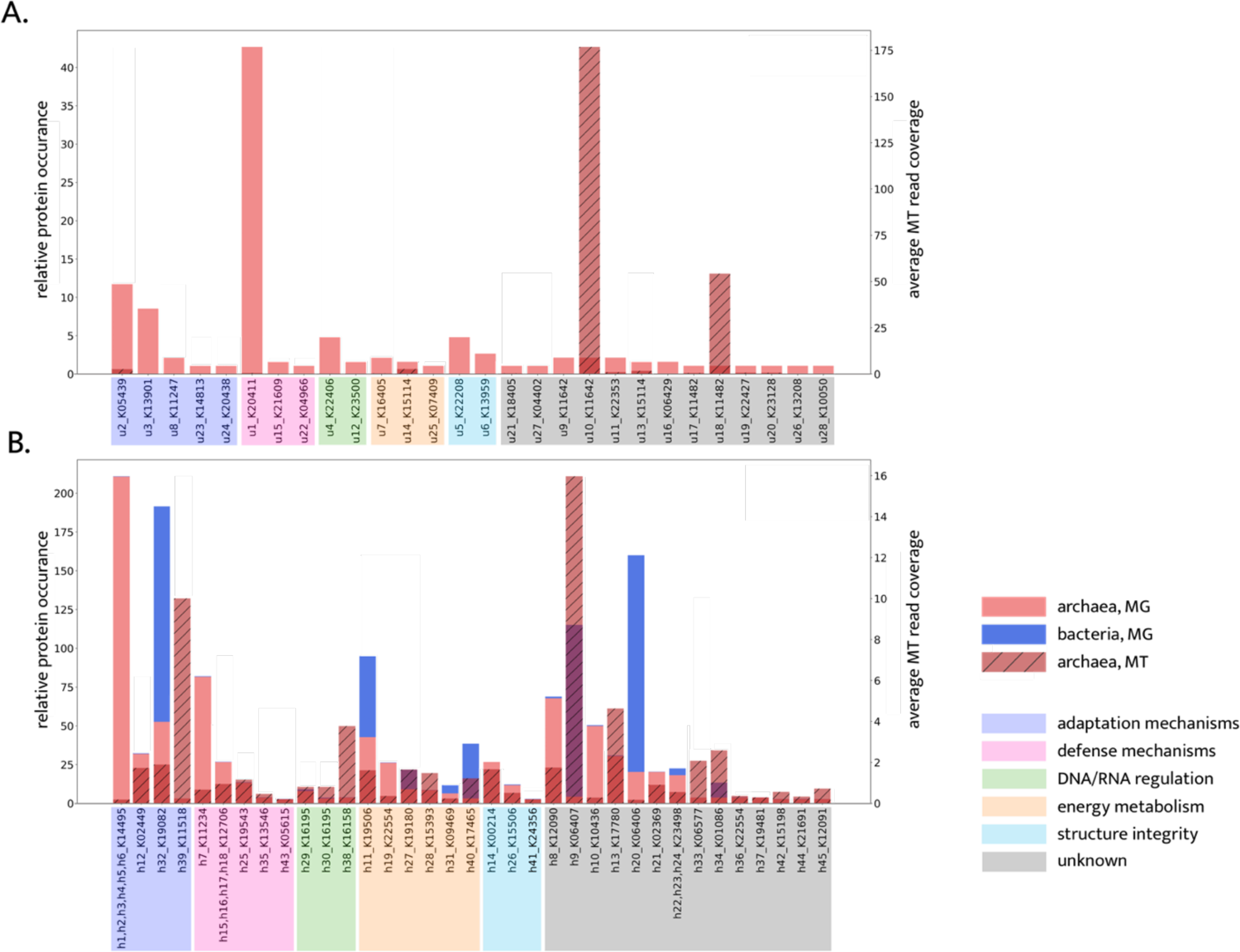
Relative occurrence and average metatranscriptomic read coverage of proteins in the **A,** unique and **B**, homologous groups of clusters with archaeal proteins. MG – metagenomics, MT – metatranscriptomics.

Our analysis demonstrated disparity in annotations between sequence- and structure-based approaches. Notably, 46% (13 out of 28) and 31% (14 out of 45) of the unique and homologous sourmash clusters, respectively, lacked structure-based annotations, suggesting a reliance on sequence information for their functional annotation thus far. However, literature searches suggest that the KEGG annotations may not provide reasonable or meaningful functional assignments for most of these unannotated proteins. For instance, a protein annotated as *mitochondrial import receptor subunit TOM40* by KEGG is predicted to be a *putative intimin/invasin-like protein* based on its structure, which is more relevant in the context of archaeal biology than being a eukaryotic protein involved in mitochondrial protein import. Similarly, a protein annotated as *Endophilin-A*, a eukaryotic protein involved in membrane curvature, shows structural similarity to *PilC, a type IVa pilus subunit* of a prokaryotic adhesion filament. While the presence of eukaryotic proteins in archaea is not surprising from an evolutionary perspective, the assignment of a protein to its evolutionary homolog from a different kingdom may not provide precise functional assignment of protein function.

In general, the agreement between the sequence- and structure-based methods was limited, with 4% (1 out of 28) and 25% (11 out of 45) of the unique and homologous proteins showing consistent annotations, respectively (Supp. Tab. 3-5). The rest of the proteins exhibited disparity between sequence- and structure-based annotations, which was assessed by comparing their reported functions. For example, unique sourmash cluster *u24* yielded different annotations using EGGNOG, KEGG, and Pfam databases which we used to potentially resolve disparities in the annotations (Supp. Tab. 3). However, a consensus structure-based annotation identified it as *polypeptide N-acetylgalactosaminyltransferase*, providing additional annotation beyond sequence analysis. Similarly, the homologous protein clusters *h15-h18* had the same functional assignments as *novobiocin biosynthesis protein NovC* using KEGG, but structure-based annotation revealed further distinctions: *h16* and *h18* were classified as members of the *LytR-Cps2A-Psr (LCP) protein family*, *h15* was annotated as *78 kDa glucose-regulated protein*, and *h17* remained unannotated (Supp. Tab. 4). The incorporation of structural information in protein annotation enables the distinction between closely related sequences, offering additional insights into protein function, which highlights the crucial role of structural data in understanding protein functionality.

We further identified glycosyltransferases responsible for N and O-linked glycosylation from clusters *h1-h6* as prevalent archaeal gut-specific proteins. These proteins may contribute to the viability and adaptability of archaeal cells in the gut. For instance, the most prevalent unique archaeal glycosyltransferase is *4-amino-4-deoxy-L-arabinose (L-Ara4N) transferase,* which is essential for the protection from environmental stress, symbiosis, virulence and resistance against antimicrobial activity [72], [73]. Moreover, one of the six glycosyltransferases is a *dolichyl-diphosphooligosaccharide – protein glycosyltransferase subunit STT3B* (*h5*) which functions as an accessory protein in N-glycosylation and provides its maximal efficiency [74]. Archaeal N-glycosylation is known to play an important role in the viability and adaptivity of archaeal cells to external conditions such as high salinity [75], elevated temperatures [76] and an acidic environment [77] while also maintaining the structural integrity of cells [78], [79]. Four out of the six identified glycosyltransferases are *dolichyl-phosphate-mannose-protein mannosyltransferases 1 (POMT1),* which are responsible for O-linked glycosylation of proteins in eukaryotes. Another O-glycosylation associated protein, *polypeptide N-acetylgalactosaminyltransferase*, was found in the subset of unique archaeal proteins (*u24*). *M. smithii* has been found to decorate its cellular surface with sugar residues mimicking those present in the glycan landscape of the intestinal environment [80]. The presence of human mucus- and epithelial cell surface-associated glycans in *M. smithii*, along with the coding potential for enzymes involved in O-linked glycosylation in archaeal gut species, suggests that *M. smithii* cells might have the capability to emulate the surfaces of eukaryotic cells in the intestinal mucus. Beyond their structural role in proteins, O-glycans can also act as regulators of protein interactions, influencing both interprotein and cell-to-cell communication processes involved in cell trafficking and environmental recognition [81].

Further findings suggest that *2-aminoethylphosphonate-pyruvate (2-AEP) aminotransferase, transthyretin-like protein* and *phosphoenolpyruvate-dependent sugar phosphotransferase system (PTS) system* encoded by *M. smithii* contribute to energy metabolism. 2-AEP is an enzyme commonly found in bacteria [82] and is known to play a critical role in phosphonate degradation [83], which serves as an important source and production pathway for methane [84].

Additionally, cold-shock domains of *Unr protein* potentially provide *M. smithii* with adaptation strategies through stress-induced control of gene expression [85]. Furthermore, the predicted involvement of proteins such as the *Specificity subunit of type I restriction-modification EcoKI enzyme* [86] and *type II restriction endonuclease BglII* [87] suggests their potential role in host defense strategies employed by *M. smithii* to protect themselves in the gut environment. Additionally, it is conceivable that archaeal proteins may play a role in protecting against toxicity from other organisms in the gut using *propanediol utilization protein pduA* [88]–[90], as well as acquiring genes of bacterial origin through horizontal gene transfer. If this is the case, the presence of adhesin-like proteins in archaea could potentially enable them to form symbiotic relationships with bacterial neighbors with diverse metabolic potentials [91]. Figure 5 provides a schematic representation emphasizing specific proteins identified in this study, which could potentially play a significant role in the functional dynamics of archaea within the human intestine. A more detailed description of all identified *M. smithii* proteins is provided in Supplementary Materials.

### Characterization of select proteins and gene structures in *Methanobrevibacter smithii* genomes

To elucidate the level of conservation among the identified genes recovered in our analyses, we assessed the level of genomic conservation within genomes of 2 strains of *M. smithii, 2 strains of Ca. Methanobreviabcter intestini and* the related species *Methanobrevibacter_A oralis* as a reference. *Ca. M. intestini* has been recently classified as an independent species within the *Methanobrevibacter smithii* clade [92]. We analysed HGT events and evaluated gene structure stability. Using 1022 available MAGs, we noted an increase in HGT events between 319 genomes of two *M. smithii* strains: *Methanobrevibacter_A smithii* and GCF_000016525.1 (based on GTDB classification) (Supp. Fig. 1). Specifically, 2.6% of the MAGs (n=27) exhibited HGT events involving the transfer of approximately 10±3 genes to other MAGs. Intriguingly, MAGs exhibiting HGT events were sampled in diverse geographical locations such as Austria, France, the UK and the US. Our results suggest that the propensity of these MAGs to exchange genomic segments may be attributed to similarities in their respective local environments [93], including dietary and lifestyle factors of the individuals. Thus, it is plausible that exposure to similar diets or stresses may have influenced the evolution of these MAGs via HGT along comparable trajectories. Conversely, the low occurrence of HGT events among the majority (97.4%) of available *M. smithii* genomes indicates their overall genomic conservation and stability. This could be explained by the fact that these MAGs were sampled from individuals living under similar dietary and lifestyle conditions. Importantly, our findings support the concept of genomic stability in *M. smithii*, as we observed a high degree of conservation in the flanking regions of the genes of interest across various *M. smithii* genomes. Through synteny analyses, we found compelling evidence of conserved synteny for genes encoded in *M. smithii* genomes (https://doi.org/10.5281/zenodo.8024791).

Among the proteins specific for gut-associated archaea, we identified *stage V sporulation proteins AE* (*spoVAE*) and AD (*spoVAD*) (*h9* and *h20*). Using BLAST searches, we extracted 250 bacterial protein sequences for *SpoVAE* and *SpoVAD* from Uniprot, including 12 *spoVAE* and 38 *spoVAD* proteins from environmental samples and the rest from isolate bacterial genomes belonging to the *Firmicutes* phylum. Phylogenetic trees demonstrated that proteins from *h9* and *h20* are phylogenetically and compositionally distinct from other sequences and form separate branches (Supp. Fig. 2-3). Gene synteny analyses revealed that sporulation genes are grouped in operons (K06405, K06406 and K06407; Fig. 3). Moreover, the flanking regions around sporulation genes include genes with key archaeal as well as methanogenic functions. In addition, the flanking regions of both *spoVAE* and *spoVAD* genes are also encoded in the *M. smithii* isolate DSM 861 genome (Fig. 4). However, in contrast to our MAGs, the isolate’s genome did not encode the *spoVAE* and *spoVAD* genes. To assess whether *spoVAE* and *spoVAD* genes were acquired by *M. smithii* via HGT, we performed synteny analysis of bacterial sequences obtained from our human gut dataset that shared similarities with the archaeal sequences in clusters *h9* and *h20*. This analysis revealed that in the bacterial genomes found in the human intestine, the flanking regions of *spoVAE* and *spoVAD* genes include genes mediating and facilitating HGT, such as a site-specific DNA recombinase (K06400) encoded upstream from *spoVAE* and type IV pilus assembly proteins (K02662, K02664) encoded downstream from *spoVAD* (Supp. Fig. 4-5). Genes originating from clusters *h9* and *h20* are found within bacterial genomes of Firmicutes phylum members, specifically *Clostridium* sp. CAG-302 and CAG-269, which highlights their association with known bacterial taxa in the gut and indicates horizontal gene transfer between these distantly related taxa.

**Figure 3.**
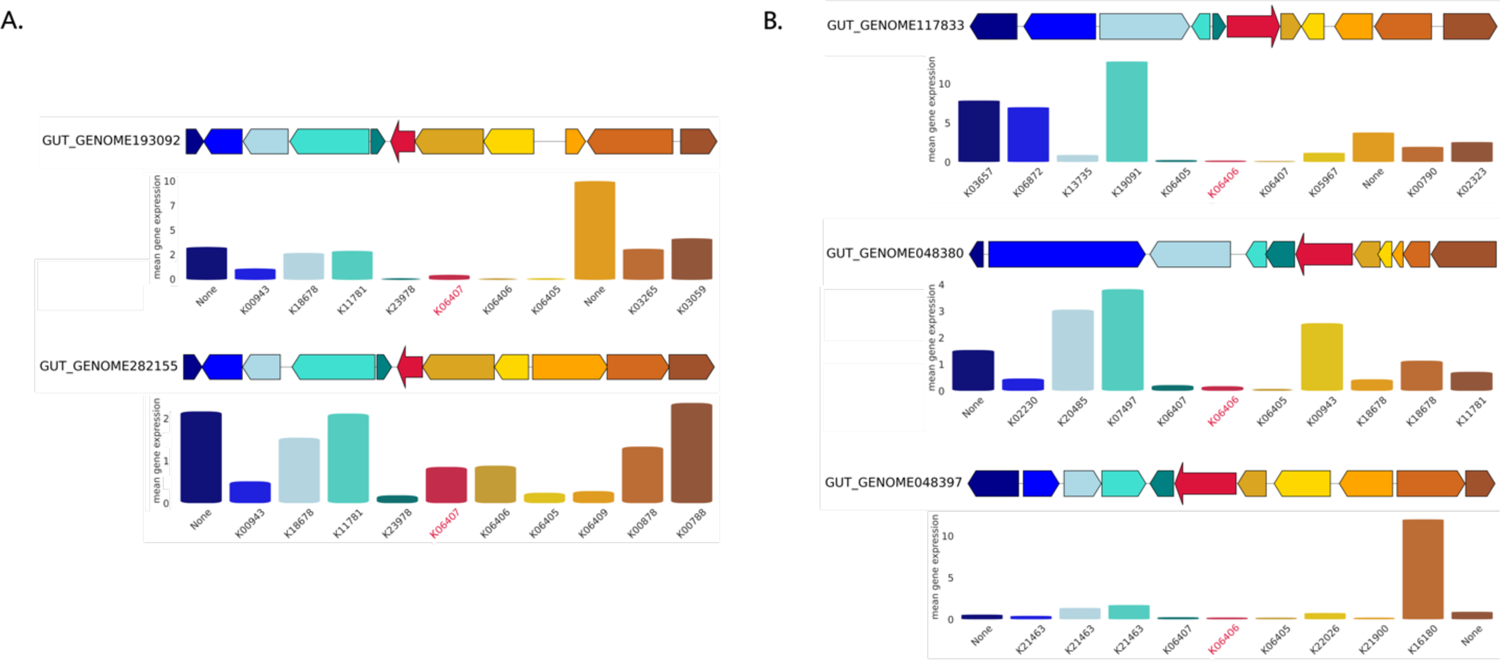
Gene synteny for sporulation stage V genes AE and AD from their respective smash clusters **A**, h9, and **B**, h20. Gene expression of target genes (spoVAE and spoVAD, in red) as well as genes from flanking regions are demonstrated below each sequence and are colored correspondingly. Genes with key archaeal functions: **A**, pyrimidine metabolism (K18678, phytol kinase), methane metabolism (K11781, 5-amino-6-(D-ribitylamino)uracil– L-tyrosine 4-hydroxyphenyl transferase) and thiamine metabolism (K00878, hydroxyethylthiazole kinase; K00788, thiamine-phosphate pyrophosphorylase); **B**, pyrimidine metabolism (K22026, nucleoside kinase; K18678, phytol kinase) and methane metabolism (K11781, 5-amino-6-(D-ribitylamino)uracil–L-tyrosine 4-hydroxyphenyl transferase).

**Figure 4.**
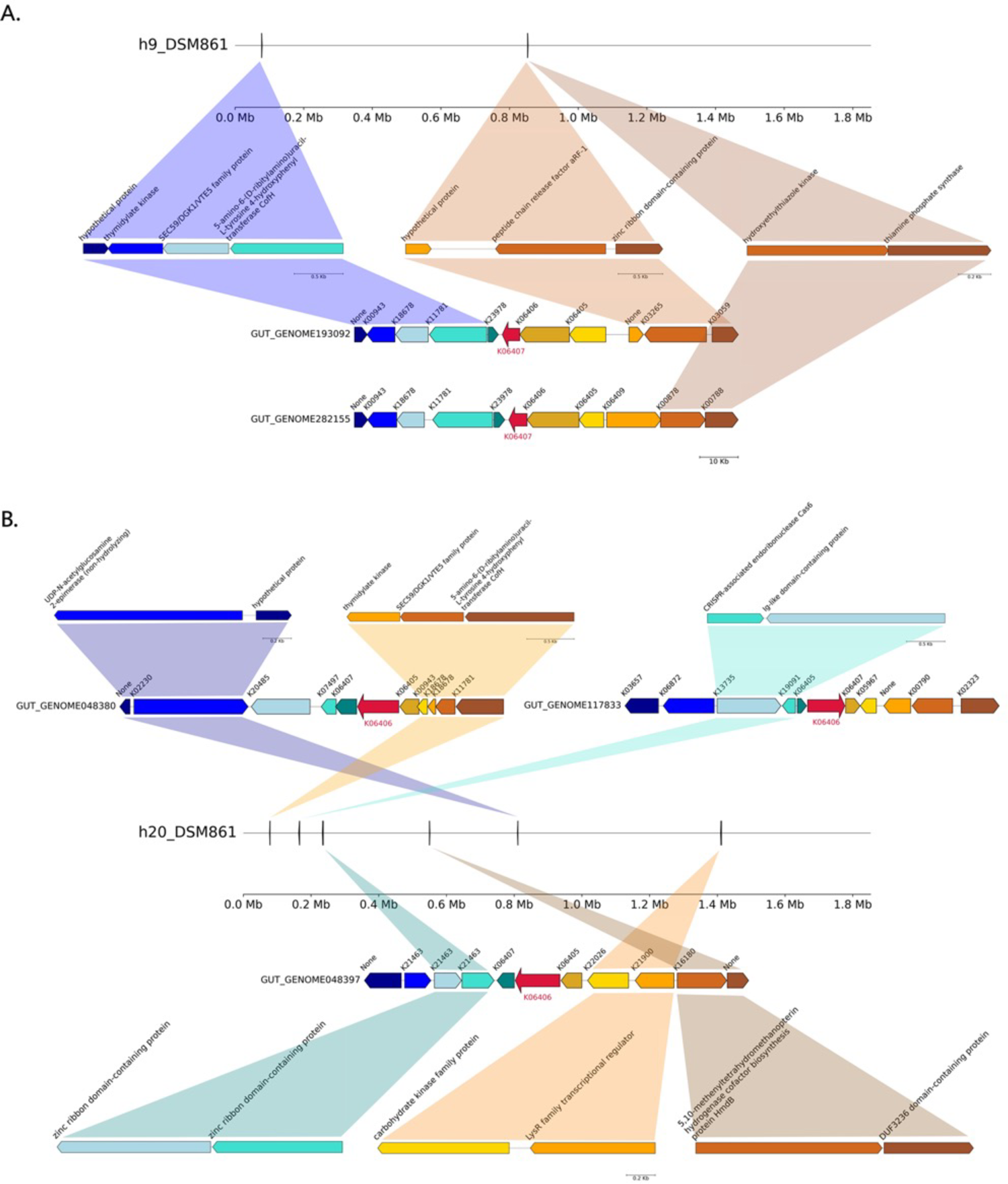
Genomic context of the archaeal flanking regions up- and downstream of the **A**, spoVAE and **B**, spoVAD gene clusters in the M. smithii strain DSM 861.

**Figure 5.**
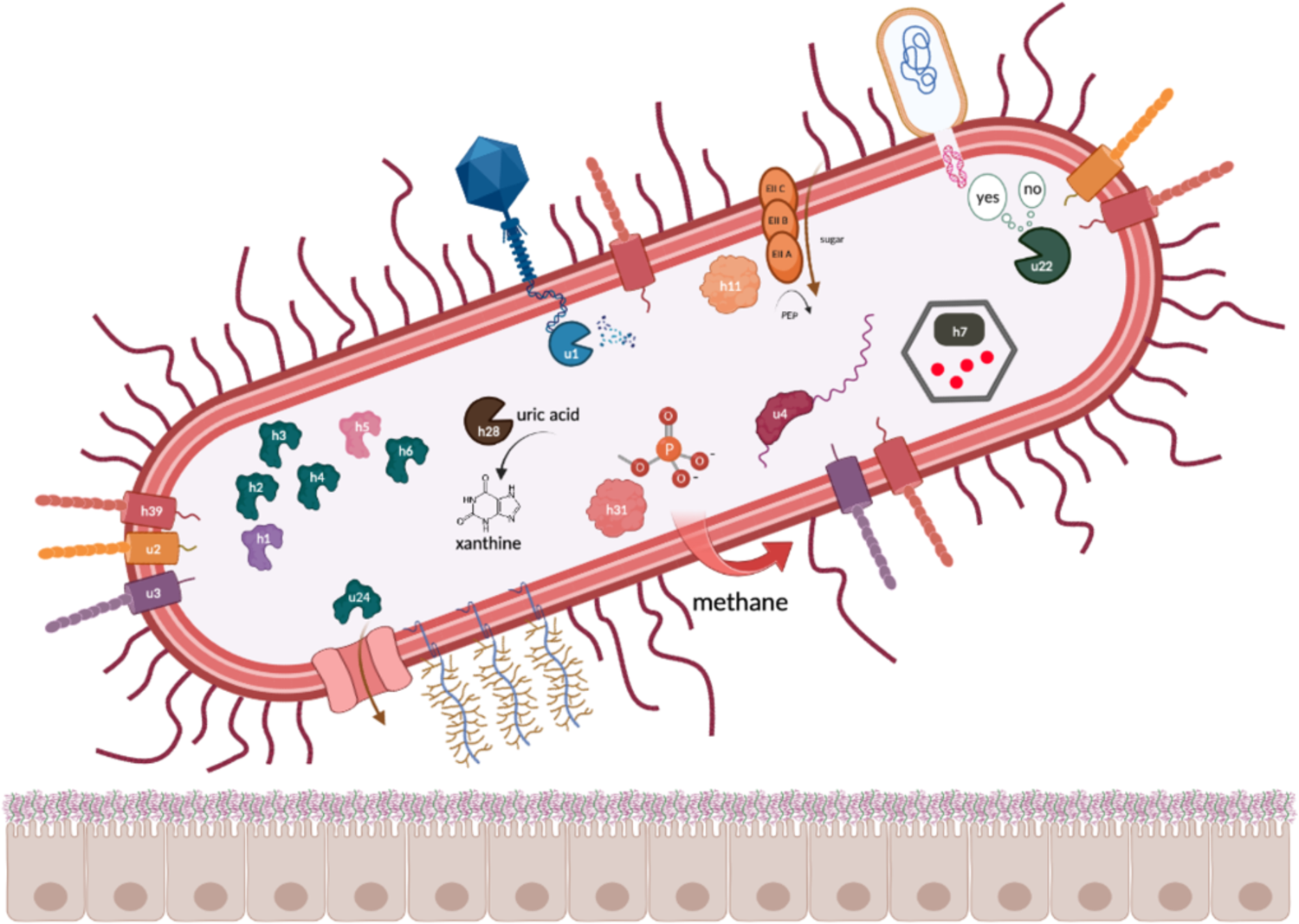
Schematic proposal highlighting proteins specific to gut-associated archaea with described functions: **u1** - Type II restriction endonuclease BglII, **u2** - Intimin/invasin-like protein with a Ig-like domain, **u3** - Intimin/invasin-like protein, **u4** - Unr protein, **u22** - Type I restriction-modification EcoKI enzyme, specificity subunit, **u24** - Polypeptide N-acetylgalactosaminyltransferase, **h1** - 4-amino-4-deoxy-L-arabinose transferase or related glycosyltransferases of PMT family, **h2,3,4,6** - Dolichyl-phosphate-mannose – protein mannosyltransferase 1, **h5** - Dolichyl-diphosphooligosaccharide-protein glycosyltransferase subunit STT3B, **h7** - Propanediol utilisation protein pduA, **h11** - Phosphoenolpyruvate-dependent PTS system, IIA component, **h28** - Transthyretin-like protein, **h31** - 2-aminoethylphosphonate-pyruvate aminotransferase.

While sporulation has been primarily observed in spore-forming bacteria and not in archaea, it is known that non-sporulating bacterial species also encode sporulation genes. In these bacterial taxa, the genes likely encode regulatory proteins involved in peptidoglycan (PPG) turnover, thereby playing a role in cell division and/or development [94], [95]. Archaea lack PPG but methanogenic archaea, including *Methanobrevibacter* species, use pseudopeptidoglycan (pseudo-PPG) instead, which functions similarly to PPG in a bacterial cell and results in Gram positive staining [96]. Certain structural similarities between methanogens and bacteria described above leave open the question of whether sporulation proteins could play a similar role in pseudopeptidoglycan turnover in methanogenic archaea, analogous to their function in non-sporulating bacteria. The identification of these genes holds significant interest, especially in light of the work by Nelson Sathi *et al.*, suggesting that methanogens frequently acquire functionally active genes through horizontal transfer from bacteria. Comprehensive experimental analysis is required to determine their specific functions, but these findings present an exciting opportunity for further exploration [97]. Phylogenetic analysis of *spoVAE* and *spoVAD* has demonstrated that sequences from the abovementioned clusters are compositionally homogeneous but phylogenetically distant from other known similar sequences in Uniprot, and therefore might be unique to the human gut environment. Moreover, archaeal and bacterial sequences from sourmash clusters *h9* and *h20* branch out together, which suggests that sporulation genes encoded in archaea are the result of horizontal gene transfer from bacteria to archaea. This study provides evidence that archaeal genomes exhibit clustered sporulation genes surrounded by genes linked to archaea-specific functions like pyrimidine, thiamine, and methane metabolism. Moreover, genes in flanking regions up- and downstream of *spoVAE* and *spoVAD* genes are indeed encoded in the representative *M. smithii* isolate DSM 861. As bacteria encoding similar *spoVAE* and *spoVAD* proteins and bacterial sequences from clusters *h9* and *h20* belong to various species of the *Clostridium* genus, HGT probably occurred in the direction from the abovementioned species to *M. smithii*. Moreover, Ruaud, Esquivel-Elizondo, de la Cuesta-Zuluaga *et al.* [98] have provided evidence of a syntrophic relationship between *Firmicutes* bacteria and *Methanobrevibacter smithii*. The co-occurrence of these microorganisms is likely facilitated by physical and metabolic interactions. In addition to this, genes *h9* and *h20* as well as their surrounding genes are expressed by the archaeal genomes sampled from human faecal samples.

## Conclusion

Our study aimed to uncover the potential functions of archaeal proteins, particularly those encoded by *M. smithii*, in the human gut. Sequence similarity-based methods, while effective for highly similar proteins (>70-80% identity), may not accurately represent the functions of archaeal proteins due to the lack of experimental validation. More specifically, publicly available databases have limited experimentally validated archaeal sequences compared to bacterial and eukaryotic proteins (∼7 000 000 archaeal, ∼166 000 000 bacterial and ∼70 000 000 eukaryotic proteins, UniProtKB Jun 2023) making sequence-based protein annotations applicable to only a subset of archaeal proteins. In contrast, recent deep learning-based methods enable protein structure prediction and annotation without relying on high sequence similarity, allowing for functional similarity beyond close sequence matches. We used structural methods to improve the annotation of archaeal proteins, gaining better insights into their functions compared to traditional sequence-based methods. This approach allowed us to refine some existing annotations and discover new functions for others, giving us valuable insights into the roles of archaeal genes in the human gut. Our findings focus on the characterization of human-associated and gut-specific proteins identified in *M. smithii,* a metabolically proficient and clinically relevant methanogenic archaeon known to be linked to gastrointestinal disorders, including IBD and obesity. Future work should help in resolving the predicted structures and protein functions using experimental approaches.

## Supporting information

Main manuscript tables

## Acknowledgments

All authors proof-read and approved of the content in this research paper. We are especially grateful for the critical feedback and suggestions provided by Dr. Cedric Laczny. Experiments presented in this work were carried out using the HPC facility of the University of Luxembourg [99]. This project has received funding from the European Research Council (ERC) under the European Union’s Horizon 2020 research and innovation program (grant agreement No. 863664).

## Competing Interests

The authors declare that they have no competing interests.

## Data Availability Statement

Microbial MAGs from UHGG collection are available from the MGnify FTP site at http://ftp.ebi.ac.uk/pub/databases/metagenomics/mgnify_genomes/, MAGs from the GEM catalog are accessible at https://portal.nersc.gov/GEM/. Metatranscriptomic sequencing reads are available from NCBI BioProject <u>PRJNA289586</u> and assembled contigs can be assessed at MG-RAST [100] (submission IDs are indicated in MT_assembly_RAST_ids.xlsx). A description of the analyses including pre-processing steps along with the scripts for the main analysis, archaeal gut-specific unique and homologous sourmash clusters and synteny plots can be found at GitLab: https://gitlab.lcsb.uni.lu/polina.novikova/archaea-in-gut.

---

**Figure 1.**
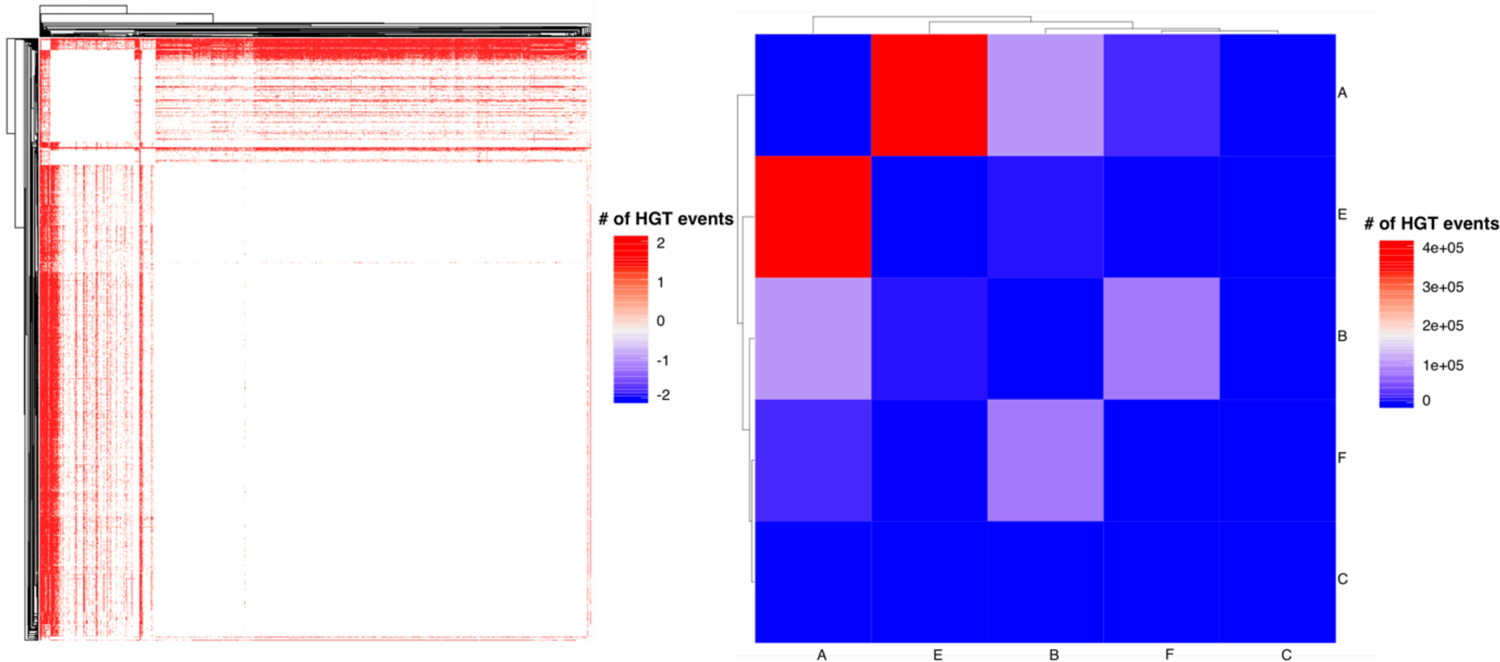
Heatmaps demonstrating the intensity of HGT events between M. smithii genomes. **A**, HGT between taxonomic groups named as follows: A - Methanobrevibacter_A smithii, B - Methanobrevibacter_A smithii_A (Ca. Methanobreviabcter intestini), C - Methanobrevibacter_A oralis, E - GCF_000016525.1 (M. smithii), F - GCF_002252585.1 (Ca. Methanobreviabcter intestini); **B**, HGT events between individual genomes of same groups. The legend depicts the frequency of HGT events among the genomes of **A**, taxonomic groups and **B**, individual genomes

**Figure 2.**
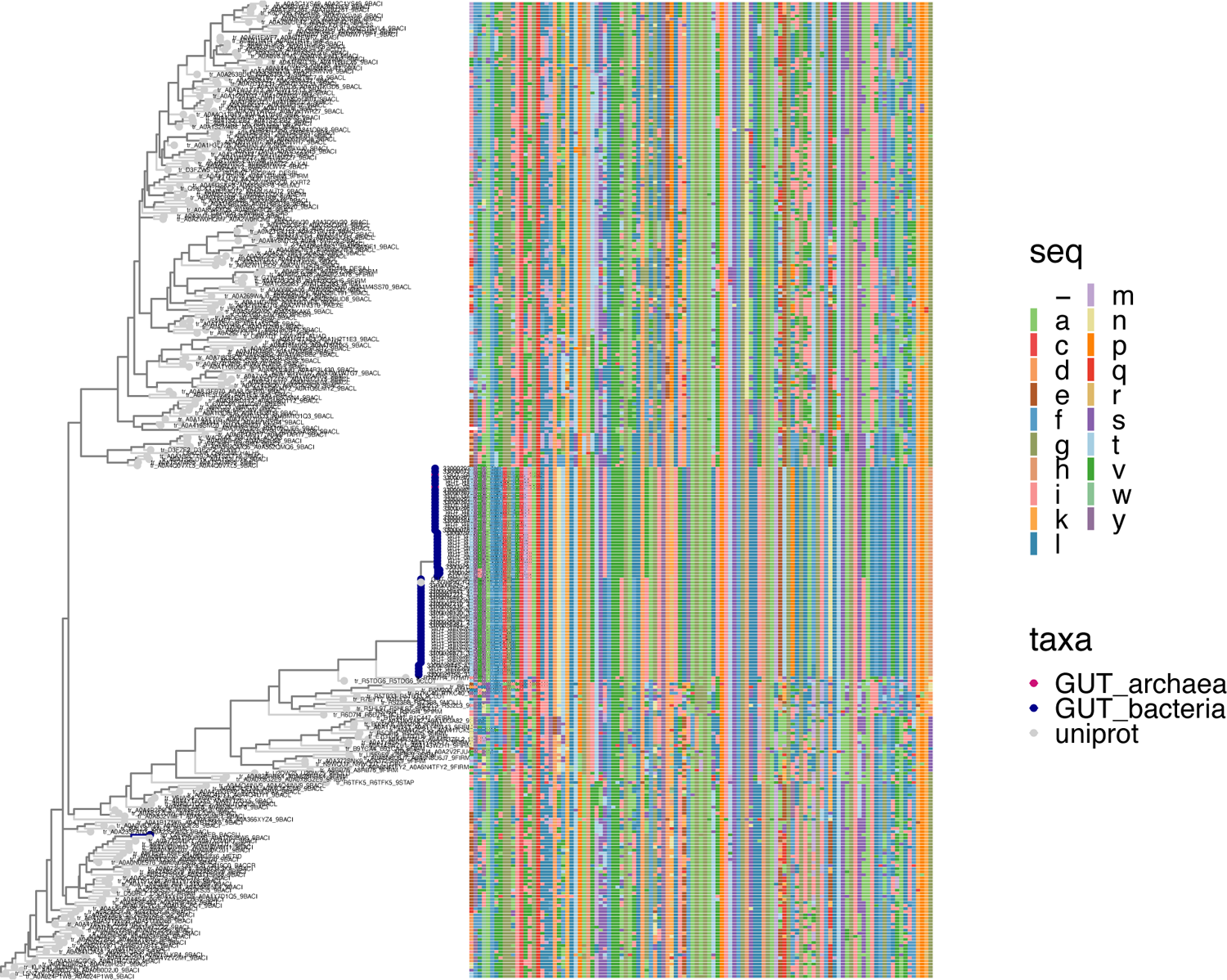
Phylogenetic tree of stage V sporulation proteins AE from identified SC h9 and Uniprot. Bacterial and archaeal proteins from cluster h9 are depicted as GUT_bacteria and GUT_archaea in dark blue and pink, respectively.

**Figure 3.**
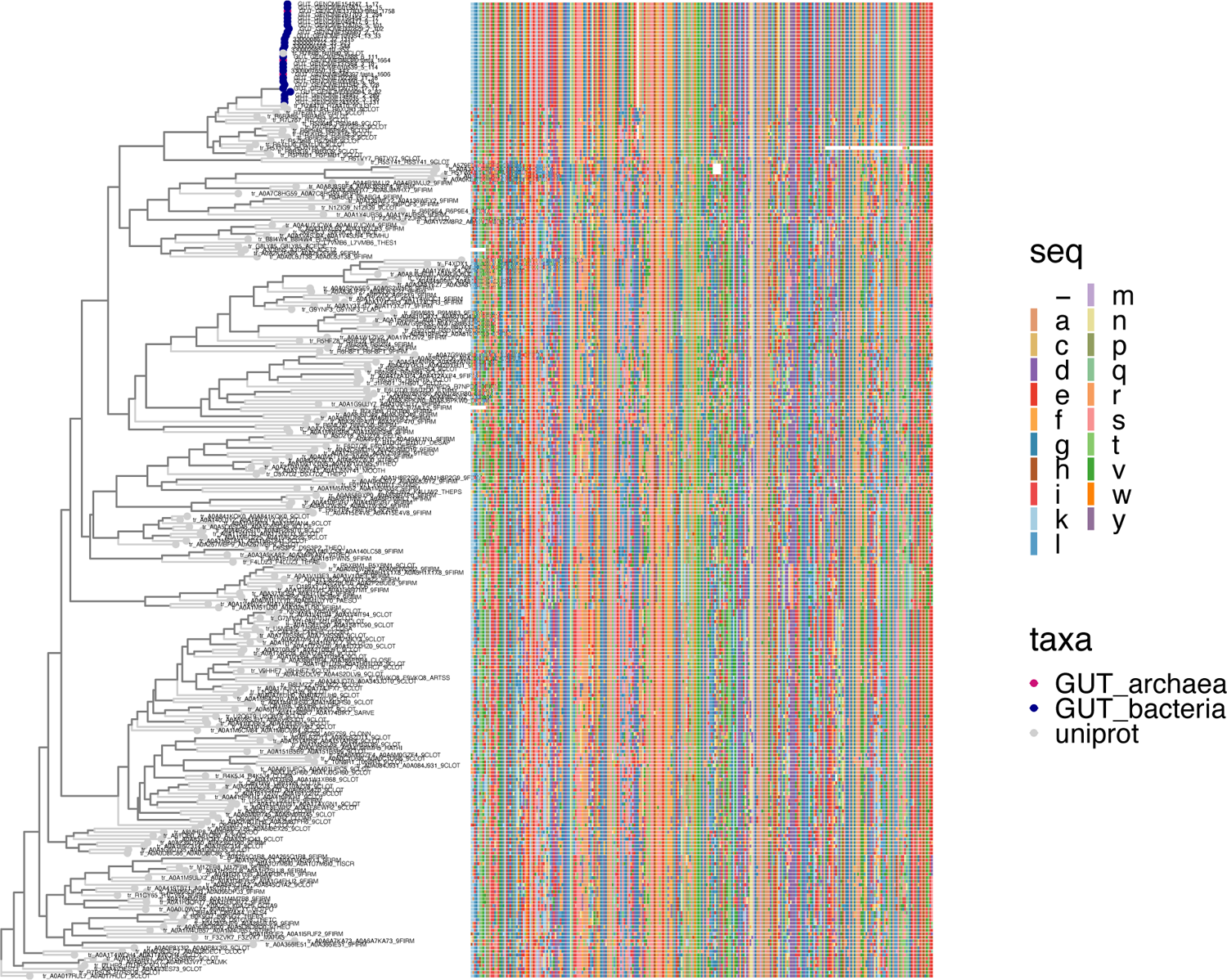
Phylogenetic tree of stage V sporulation proteins AD from identified SC h20 and Uniprot. Bacterial and archaeal proteins from cluster h9 are depicted as GUT_bacteria and GUT_archaea in dark blue and pink, respectively.

**Figure 4.**
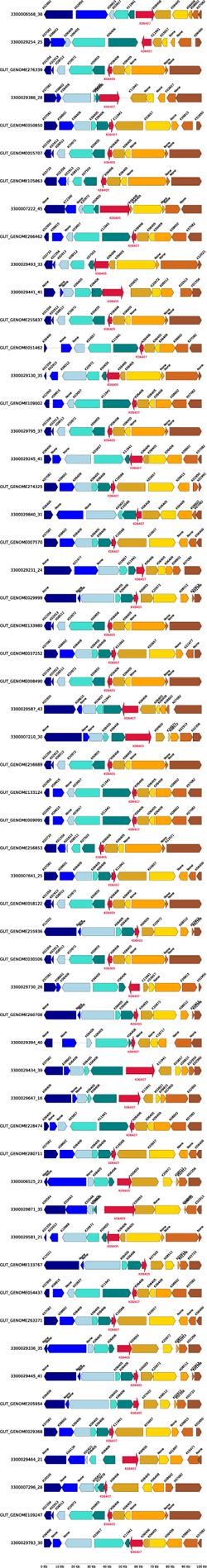
Gene synteny of homologous bacterial sequences obtained from the human gut dataset that share similarities with the archaeal sequences from cluster h9 encoding stage V sporulation protein AE (spoVAE)

**Figure 5.**
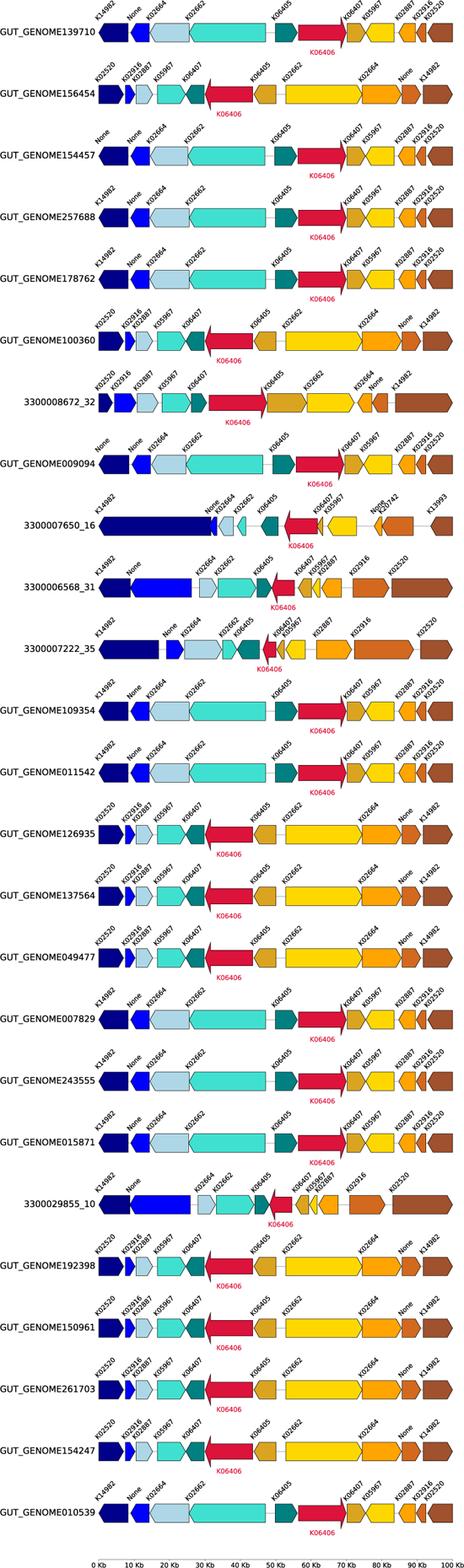
Gene synteny of homologous bacterial sequences obtained from the human gut dataset that share similarities with the archaeal sequences from cluster h20 encoding stage V sporulation protein AD (spoVAD).

## Notes

### Competing Interest Statement

The authors have declared no competing interest.

### Summary of Updates

The text is revised and rearranged. Supplemental files and table are updated.

## References

[1] C. R. Woese and G. E. Fox, ‘Phylogenetic structure of the prokaryotic domain: The primary kingdoms’, Proc. Natl. Acad. Sci., vol. 74, no. 11, pp. 5088–5090, Nov. 1977, doi: 10.1073/pnas.74.11.5088.

[2] W. E. Balch, L. J. Magrum, G. E. Fox, R. S. Wolfe, and C. R. Woese, ‘An ancient divergence among the bacteria’, J. Mol. Evol., vol. 9, no. 4, pp. 305–311, Aug. 1977, doi: 10.1007/BF01796092.

[3] G. E. Fox, L. J. Magrum, W. E. Balch, R. S. Wolfe, and C. R. Woese, ‘Classification of methanogenic bacteria by 16S ribosomal RNA characterization’, Proc. Natl. Acad. Sci. U. S. A., vol. 74, no. 10, pp. 4537–4541, Oct. 1977, doi: 10.1073/pnas.74.10.4537.

[4] C. R. Woese, O. Kandler, and M. L. Wheelis, ‘Towards a natural system of organisms: proposal for the domains Archaea, Bacteria, and Eucarya’, Proc. Natl. Acad. Sci. U. S. A., vol. 87, no. 12, pp. 4576–4579, Jun. 1990, doi: 10.1073/pnas.87.12.4576.

[5] Y. Liu et al., ‘Expanded diversity of Asgard archaea and their relationships with eukaryotes’, Nature, vol. 593, no. 7860, Art. no. 7860, May 2021, doi: 10.1038/s41586-021-03494-3.

[6] T. A. Williams, C. J. Cox, P. G. Foster, G. J. Szöllősi, and T. M. Embley, ‘Phylogenomics provides robust support for a two-domains tree of life’, *Nat*. Ecol. Evol., vol. 4, no. 1, pp. 138–147, Jan. 2020, doi: 10.1038/s41559-019-1040-x.

[7] M. Könneke, A. E. Bernhard, J. R. de la Torre, C. B. Walker, J. B. Waterbury, and D. A. Stahl, ‘Isolation of an autotrophic ammonia-oxidizing marine archaeon’, Nature, vol. 437, no. 7058, pp. 543–546, Sep. 2005, doi: 10.1038/nature03911.

[8] M. Pester, C. Schleper, and M. Wagner, ‘The Thaumarchaeota: an emerging view of their phylogeny and ecophysiology’, Curr. Opin. Microbiol., vol. 14, no. 3, pp. 300–306, Jun. 2011, doi: 10.1016/j.mib.2011.04.007.

[9] E. F. DeLong, ‘Everything in moderation: archaea as “non-extremophiles”’, Curr. Opin. Genet. Dev., vol. 8, no. 6, pp. 649–654, Dec. 1998, doi: 10.1016/s0959-437x(98)80032-4.

[10] C. Schleper, ‘Ammonia oxidation: different niches for bacteria and archaea?’, ISME J., vol. 4, no. 9, pp. 1092– 1094, Sep. 2010, doi: 10.1038/ismej.2010.111.

[11] D. L. Valentine, ‘Adaptations to energy stress dictate the ecology and evolution of the Archaea’, Nat. Rev. Microbiol., vol. 5, no. 4, pp. 316–323, Apr. 2007, doi: 10.1038/nrmicro1619.

[12] C. Hoegenauer, H. F. Hammer, A. Mahnert, and C. Moissl-Eichinger, ‘Methanogenic archaea in the human gastrointestinal tract’, Nat. Rev. Gastroenterol. Hepatol., vol. 19, no. 12, Art. no. 12, Dec. 2022, doi: 10.1038/s41575-022-00673-z.

[13] C. M. Thomas, E. Desmond-Le Quéméner, S. Gribaldo, and G. Borrel, ‘Factors shaping the abundance and diversity of the gut archaeome across the animal kingdom’, Nat. Commun., vol. 13, no. 1, Art. no. 1, Jun. 2022, doi: 10.1038/s41467-022-31038-4.

[14] C. Moissl-Eichinger et al., ‘Human age and skin physiology shape diversity and abundance of Archaea on skin’, Sci. Rep., vol. 7, no. 1, p. 4039, Jun. 2017, doi: 10.1038/s41598-017-04197-4.

[15] A. J. Probst, A. K. Auerbach, and C. Moissl-Eichinger, ‘Archaea on Human Skin’, PLOS ONE, vol. 8, no. 6, p. e65388, Jun. 2013, doi: 10.1371/journal.pone.0065388.

[16] C. Kumpitsch, K. Koskinen, V. Schöpf, and C. Moissl-Eichinger, ‘The microbiome of the upper respiratory tract in health and disease’, BMC Biol., vol. 17, no. 1, p. 87, Nov. 2019, doi: 10.1186/s12915-019-0703-z.

[17] E. Sogodogo et al., ‘First characterization of methanogens in oral cavity in Malian patients with oral cavity pathologies’, BMC Oral Health, vol. 19, no. 1, p. 232, Oct. 2019, doi: 10.1186/s12903-019-0929-8.

[18] J. Y. Kim et al., ‘The human gut archaeome: identification of diverse haloarchaea in Korean subjects’, Microbiome, vol. 8, no. 1, p. 114, Aug. 2020, doi: 10.1186/s40168-020-00894-x.

[19] P. B. Eckburg et al., ‘Diversity of the human intestinal microbial flora’, Science, vol. 308, no. 5728, pp. 1635–1638, Jun. 2005, doi: 10.1126/science.1110591.

[20] S. B. Ghavami et al., ‘Alterations of the human gut Methanobrevibacter smithii as a biomarker for inflammatory bowel diseases’, Microb. Pathog., vol. 117, pp. 285–289, Apr. 2018, doi: 10.1016/j.micpath.2018.01.029.

[21] Y. Houshyar, L. Massimino, L. A. Lamparelli, S. Danese, and F. Ungaro, ‘Going Beyond Bacteria: Uncovering the Role of Archaeome and Mycobiome in Inflammatory Bowel Disease’, Front. Physiol., vol. 12, p. 783295, Dec. 2021, doi: 10.3389/fphys.2021.783295.

[22] R. J. Basseri et al., ‘Intestinal methane production in obese individuals is associated with a higher body mass index’, Gastroenterol. Hepatol., vol. 8, no. 1, pp. 22–28, Jan. 2012.

[23] B. S. Samuel and J. I. Gordon, ‘A humanized gnotobiotic mouse model of host–archaeal–bacterial mutualism’, Proc. Natl. Acad. Sci., vol. 103, no. 26, pp. 10011–10016, Jun. 2006, doi: 10.1073/pnas.0602187103.

[24] R. Mathur, M. Amichai, K. S. Chua, J. Mirocha, G. M. Barlow, and M. Pimentel, ‘Methane and Hydrogen Positivity on Breath Test Is Associated With Greater Body Mass Index and Body Fat’, J. Clin. Endocrinol. Metab., vol. 98, no. 4, pp. E698–E702, Apr. 2013, doi: 10.1210/jc.2012-3144.

[25] G. Borrel, J.-F. Brugère, S. Gribaldo, R. A. Schmitz, and C. Moissl-Eichinger, ‘The host-associated archaeome’, Nat. Rev. Microbiol., vol. 18, no. 11, Art. no. 11, Nov. 2020, doi: 10.1038/s41579-020-0407-y.

[26] G. Borrel et al., ‘Genomics and metagenomics of trimethylamine-utilizing Archaea in the human gut microbiome’, ISME J., vol. 11, no. 9, pp. 2059–2074, Sep. 2017, doi: 10.1038/ismej.2017.72.

[27] C. Bang, K. Weidenbach, T. Gutsmann, H. Heine, and R. A. Schmitz, ‘The Intestinal Archaea Methanosphaera stadtmanae and Methanobrevibacter smithii Activate Human Dendritic Cells’, PLoS ONE, vol. 9, no. 6, p. e99411, Jun. 2014, doi: 10.1371/journal.pone.0099411.

[28] Z. Lyu and W. B. Whitman, ‘Transplanting the pathway engineering toolbox to methanogens’, Curr. Opin. Biotechnol., vol. 59, pp. 46–54, Oct. 2019, doi: 10.1016/j.copbio.2019.02.009.

[29] J. Thomsen, K. Weidenbach, W. W. Metcalf, and R. A. Schmitz, ‘Genetic Methods and Construction of Chromosomal Mutations in Methanogenic Archaea’, Methods Mol. Biol. Clifton NJ, vol. 2522, pp. 105–117, 2022, doi: 10.1007/978-1-0716-2445-6_6.

[30] A. Tebbe et al., ‘Analysis of the cytosolic proteome of Halobacterium salinarum and its implication for genome annotation’, Proteomics, vol. 5, no. 1, pp. 168–179, Jan. 2005, doi: 10.1002/pmic.200400910.

[31] K. Zaremba-Niedzwiedzka et al., ‘Asgard archaea illuminate the origin of eukaryotic cellular complexity’, Nature, vol. 541, no. 7637, pp. 353–358, Jan. 2017, doi: 10.1038/nature21031.

[32] A. Spang et al., ‘Complex archaea that bridge the gap between prokaryotes and eukaryotes’, Nature, vol. 521, no. 7551, pp. 173–179, May 2015, doi: 10.1038/nature14447.

[33] C. J. Castelle et al., ‘Genomic Expansion of Domain Archaea Highlights Roles for Organisms from New Phyla in Anaerobic Carbon Cycling’, Curr. Biol., vol. 25, no. 6, pp. 690–701, Mar. 2015, doi: 10.1016/j.cub.2015.01.014.

[34] P. Bork, ‘Powers and Pitfalls in Sequence Analysis: The 70% Hurdle’, Genome Res., vol. 10, no. 4, pp. 398–400, Jan. 2000, doi: 10.1101/gr.10.4.398.

[35] J. C. Wooley, A. Godzik, and I. Friedberg, ‘A Primer on Metagenomics’, PLOS Comput. Biol., vol. 6, no. 2, p. e1000667, Feb. 2010, doi: 10.1371/journal.pcbi.1000667.

[36] K. W. Ellens, N. Christian, C. Singh, V. P. Satagopam, P. May, and C. L. Linster, ‘Confronting the catalytic dark matter encoded by sequenced genomes’, Nucleic Acids Res., vol. 45, no. 20, pp. 11495–11514, Nov. 2017, doi: 10.1093/nar/gkx937.

[37] K. S. Makarova, Y. I. Wolf, and E. V. Koonin, ‘Towards functional characterization of archaeal genomic dark matter’, Biochem. Soc. Trans., vol. 47, no. 1, pp. 389–398, Feb. 2019, doi: 10.1042/BST20180560.

[38] V. Márquez et al., ‘Proteomic characterization of archaeal ribosomes reveals the presence of novel archaeal-specific ribosomal proteins’, J. Mol. Biol., vol. 405, no. 5, pp. 1215–1232, Feb. 2011, doi: 10.1016/j.jmb.2010.11.055.

[39] P. Wu, J. S. Brockenbrough, M. R. Paddy, and J. P. Aris, ‘NCL1, a novel gene for a non-essential nuclear protein in Saccharomyces cerevisiae’, Gene, vol. 220, pp. 109–117, Oct. 1998, doi: 10.1016/s0378-1119(98)00330-8.

[40] N. Vakirlis, A.-R. Carvunis, and A. McLysaght, ‘Synteny-based analyses indicate that sequence divergence is not the main source of orphan genes’, eLife, vol. 9, 2020, doi: 10.7554/eLife.53500.

[41] C. M. Weisman, A. W. Murray, and S. R. Eddy, ‘Many, but not all, lineage-specific genes can be explained by homology detection failure’, PLoS Biol., vol. 18, no. 11, p. e3000862, Nov. 2020, doi: 10.1371/journal.pbio.3000862.

[42] S. Sinha, A. M. Lynn, and D. K. Desai, ‘Implementation of homology based and non-homology based computational methods for the identification and annotation of orphan enzymes: using Mycobacterium tuberculosis H37Rv as a case study’, BMC Bioinformatics, vol. 21, no. 1, p. 466, Oct. 2020, doi: 10.1186/s12859-020-03794-x.

[43] A. Mahnert, M. Blohs, M.-R. Pausan, and C. Moissl-Eichinger, ‘The human archaeome: methodological pitfalls and knowledge gaps’, Emerg. Top. Life Sci., vol. 2, no. 4, pp. 469–482, Dec. 2018, doi: 10.1042/ETLS20180037.

[44] J. D. Watson et al., ‘Towards Fully Automated Structure-Based Function Prediction In Structural Genomics: A Case Study’, J. Mol. Biol., vol. 367, no. 5, pp. 1511–1522, Apr. 2007, doi: 10.1016/j.jmb.2007.01.063.

[45] C. Chothia and A. M. Lesk, ‘The relation between the divergence of sequence and structure in proteins’, EMBO J., vol. 5, no. 4, pp. 823–826, Apr. 1986, doi: 10.1002/j.1460-2075.1986.tb04288.x.

[46] J. Skolnick, M. Gao, H. Zhou, and S. Singh, ‘AlphaFold 2: Why It Works and Its Implications for Understanding the Relationships of Protein Sequence, Structure, and Function’, J. Chem. Inf. Model., vol. 61, no. 10, pp. 4827– 4831, Oct. 2021, doi: 10.1021/acs.jcim.1c01114.

[47] J. Jumper et al., ‘Highly accurate protein structure prediction with AlphaFold’, Nature, vol. 596, no. 7873, pp. 583– 589, Aug. 2021, doi: 10.1038/s41586-021-03819-2.

[48] Z. Du et al., ‘The trRosetta server for fast and accurate protein structure prediction’, Nat. Protoc., vol. 16, no. 12, pp. 5634–5651, Dec. 2021, doi: 10.1038/s41596-021-00628-9.

[49] S. Nayfach et al., ‘A genomic catalog of Earth’s microbiomes’, Nat. Biotechnol., vol. 39, no. 4, pp. 499–509, Apr. 2021, doi: 10.1038/s41587-020-0718-6.

[50] A. Almeida et al., ‘A unified catalog of 204,938 reference genomes from the human gut microbiome’, Nat. Biotechnol., vol. 39, no. 1, pp. 105–114, Jan. 2021, doi: 10.1038/s41587-020-0603-3.

[51] ‘Prodigal: prokaryotic gene recognition and translation initiation site identification | BMC Bioinformatics | Full Text’. https://bmcbioinformatics.biomedcentral.com/articles/10.1186/1471-2105-11-119 (accessed Jan. 19, 2022).

[52] M. Steinegger and J. Söding, ‘MMseqs2 enables sensitive protein sequence searching for the analysis of massive data sets’, Nat. Biotechnol., vol. 35, no. 11, pp. 1026–1028, Nov. 2017, doi: 10.1038/nbt.3988.

[53] M. Steinegger and J. Söding, ‘Clustering huge protein sequence sets in linear time’, Nat. Commun., vol. 9, no. 1, p. 2542, Jun. 2018, doi: 10.1038/s41467-018-04964-5.

[54] C. T. Brown and L. Irber, ‘sourmash: a library for MinHash sketching of DNA’, J. Open Source Softw., vol. 1, no. 5, p. 27, Sep. 2016, doi: 10.21105/joss.00027.

[55] N. T. Pierce, L. Irber, T. Reiter, P. Brooks, and C. T. Brown, ‘Large-scale sequence comparisons with sourmash’, F1000Research, vol. 8, p. 1006, Jul. 2019, doi: 10.12688/f1000research.19675.1.

[56] P. Queirós, F. Delogu, O. Hickl, P. May, and P. Wilmes, ‘Mantis: flexible and consensus-driven genome annotation’, GigaScience, vol. 10, no. 6, p. giab042, Jun. 2021, doi: 10.1093/gigascience/giab042.

[57] Moult, J., Fidelis, K., Kryshtafovych, A., Schwede, T. & Topf, M., ‘Critical assessment of techniques for protein structure prediction, fourteenth round.’, ASP 14 Abstract Book, 2020, [Online]. Available: https://www.predictioncenter.org/casp14/doc/CASP14_Abstracts.pdf

[58] R. A. Laskowski, J. D. Watson, and J. M. Thornton, ‘ProFunc: a server for predicting protein function from 3D structure’, Nucleic Acids Res., vol. 33, no. Web Server issue, pp. W89-93, Jul. 2005, doi: 10.1093/nar/gki414.

[59] V. Gligorijević et al., ‘Structure-based protein function prediction using graph convolutional networks’, Nat. Commun., vol. 12, no. 1, p. 3168, May 2021, doi: 10.1038/s41467-021-23303-9.

[60] A. Heintz-Buschart et al., ‘Integrated multi-omics of the human gut microbiome in a case study of familial type 1 diabetes’, Nat. Microbiol., vol. 2, no. 1, Art. no. 1, Oct. 2016, doi: 10.1038/nmicrobiol.2016.180.

[61] H. Li, ‘Aligning sequence reads, clone sequences and assembly contigs with BWA-MEM’, 2013, doi: 10.6084/M9.FIGSHARE.963153.V1.

[62] H. Li et al., ‘The Sequence Alignment/Map format and SAMtools’, Bioinformatics, vol. 25, no. 16, pp. 2078–2079, Aug. 2009, doi: 10.1093/bioinformatics/btp352.

[63] B. S. Pedersen and A. R. Quinlan, ‘Mosdepth: quick coverage calculation for genomes and exomes’, Bioinforma. Oxf. Engl., vol. 34, no. 5, pp. 867–868, Mar. 2018, doi: 10.1093/bioinformatics/btx699.

[64] W. Song, B. Wemheuer, S. Zhang, K. Steensen, and T. Thomas, ‘MetaCHIP: community-level horizontal gene transfer identification through the combination of best-match and phylogenetic approaches’, Microbiome, vol. 7, no. 1, p. 36, Mar. 2019, doi: 10.1186/s40168-019-0649-y.

[65] Y. Shimoyama, ‘pyGenomeViz: A genome visualization python package for comparative genomics’. Jun. 2022. Accessed: Jul. 23, 2022. [Online]. Available: https://github.com/moshi4/pyGenomeViz

[66] R. Apweiler et al., ‘UniProt: the Universal Protein knowledgebase’, Nucleic Acids Res., vol. 32, no. Database issue, pp. D115-119, Jan. 2004, doi: 10.1093/nar/gkh131.

[67] ‘BLAST: Basic Local Alignment Search Tool’. https://blast.ncbi.nlm.nih.gov/Blast.cgi (accessed Dec. 06, 2022).

[68] K. Katoh, K. Misawa, K. Kuma, and T. Miyata, ‘MAFFT: a novel method for rapid multiple sequence alignment based on fast Fourier transform’, Nucleic Acids Res., vol. 30, no. 14, pp. 3059–3066, Jul. 2002, doi: 10.1093/nar/gkf436.

[69] A. Criscuolo and S. Gribaldo, ‘BMGE (Block Mapping and Gathering with Entropy): a new software for selection of phylogenetic informative regions from multiple sequence alignments’, BMC Evol. Biol., vol. 10, p. 210, Jul. 2010, doi: 10.1186/1471-2148-10-210.

[70] L.-T. Nguyen, H. A. Schmidt, A. von Haeseler, and B. Q. Minh, ‘IQ-TREE: A Fast and Effective Stochastic Algorithm for Estimating Maximum-Likelihood Phylogenies’, Mol. Biol. Evol., vol. 32, no. 1, pp. 268–274, Jan. 2015, doi: 10.1093/molbev/msu300.

[71] G. Yu, ‘Using ggtree to Visualize Data on Tree-Like Structures’, Curr. Protoc. Bioinforma., vol. 69, no. 1, p. e96, Mar. 2020, doi: 10.1002/cpbi.96.

[72] A. Anandan and A. Vrielink, ‘Structure and function of lipid A-modifying enzymes’, Ann. N. Y. Acad. Sci., vol. 1459, no. 1, pp. 19–37, Jan. 2020, doi: 10.1111/nyas.14244.

[73] S. D. Breazeale, A. A. Ribeiro, and C. R. H. Raetz, ‘Origin of Lipid A Species Modified with 4-Amino-4-deoxy-l-arabinose in Polymyxin-resistant Mutants of Escherichia coli: AN AMINOTRANSFERASE (ArnB) THAT GENERATES UDP-4-AMINO-4-DEOXY-l-ARABINOSE *’, J. Biol. Chem., vol. 278, no. 27, pp. 24731–24739, Jul. 2003, doi: 10.1074/jbc.M304043200.

[74] A. Dell, A. Galadari, F. Sastre, and P. Hitchen, ‘Similarities and Differences in the Glycosylation Mechanisms in Prokaryotes and Eukaryotes’, Int. J. Microbiol., vol. 2010, p. e148178, Jan. 2011, doi: 10.1155/2010/148178.

[75] M. Abu-Qarn and J. Eichler, ‘Protein N-glycosylation in Archaea: defining Haloferax volcanii genes involved in S-layer glycoprotein glycosylation’, Mol. Microbiol., vol. 61, no. 2, pp. 511–525, Jul. 2006, doi: 10.1111/j.1365-2958.2006.05252.x.

[76] U. Kärcher et al., ‘Primary structure of the heterosaccharide of the surface glycoprotein of Methanothermus fervidus.’, J. Biol. Chem., vol. 268, no. 36, pp. 26821–26826, Dec. 1993, doi: 10.1016/S0021-9258(19)74185-4.

[77] U. Zähringer, H. Moll, T. Hettmann, Y. A. Knirel, and G. Schäfer, ‘Cytochrome b558/566 from the archaeon Sulfolobus acidocaldarius has a unique Asn-linked highly branched hexasaccharide chain containing 6-sulfoquinovose’, Eur. J. Biochem., vol. 267, no. 13, pp. 4144–4149, Jul. 2000, doi: 10.1046/j.1432-1327.2000.01446.x.

[78] M. F. Mescher and J. L. Strominger, ‘Purification and characterization of a prokaryotic glycoprotein from the cell envelope of Halobacterium salinarium’, J. Biol. Chem., vol. 251, no. 7, pp. 2005–2014, Apr. 1976.

[79] A. Tamir and J. Eichler, ‘N-Glycosylation Is Important for Proper Haloferax volcanii S-Layer Stability and Function’, Appl. Environ. Microbiol., vol. 83, no. 6, pp. e03152-16, Mar. 2017, doi: 10.1128/AEM.03152-16.

[80] B. S. Samuel et al., ‘Genomic and metabolic adaptations of Methanobrevibacter smithii to the human gut’, Proc. Natl. Acad. Sci. U. S. A., vol. 104, no. 25, pp. 10643–10648, Jun. 2007, doi: 10.1073/pnas.0704189104.

[81] H. H. Wandall, M. A. I. Nielsen, S. King-Smith, N. de Haan, and I. Bagdonaite, ‘Global functions of O-glycosylation: promises and challenges in O-glycobiology’, FEBS J., vol. 288, no. 24, pp. 7183–7212, 2021, doi: 10.1111/febs.16148.

[82] W. W. Metcalf and W. A. van der Donk, ‘Biosynthesis of Phosphonic and Phosphinic Acid Natural Products’, Annu. Rev. Biochem., vol. 78, pp. 65–94, 2009, doi: 10.1146/annurev.biochem.78.091707.100215.

[83] E. Zangelmi, T. Stanković, M. Malatesta, D. Acquotti, K. Pallitsch, and A. Peracchi, ‘Discovery of a New, Recurrent Enzyme in Bacterial Phosphonate Degradation: (R)-1-Hydroxy-2-aminoethylphosphonate Ammonia-lyase’, Biochemistry, vol. 60, no. 15, pp. 1214–1225, Apr. 2021, doi: 10.1021/acs.biochem.1c00092.

[84] W. W. Metcalf et al., ‘Synthesis of Methylphosphonic Acid by Marine Microbes: A Source for Methane in the Aerobic Ocean’, Science, vol. 337, no. 6098, pp. 1104–1107, Aug. 2012, doi: 10.1126/science.1219875.

[85] V. Dormoy-Raclet et al., ‘Unr, a cytoplasmic RNA-binding protein with cold-shock domains, is involved in control of apoptosis in ES and HuH7 cells’, Oncogene, vol. 26, no. 18, pp. 2595–2605, Apr. 2007, doi: 10.1038/sj.onc.1210068.

[86] L. Roer, F. M. Aarestrup, and H. Hasman, ‘The EcoKI Type I Restriction-Modification System in Escherichia coli Affects but Is Not an Absolute Barrier for Conjugation’, J. Bacteriol., vol. 197, no. 2, pp. 337–342, Jan. 2015, doi: 10.1128/JB.02418-14.

[87] A. Pingoud, M. Fuxreiter, V. Pingoud, and W. Wende, ‘Type II restriction endonucleases: structure and mechanism’, Cell. Mol. Life Sci. CMLS, vol. 62, no. 6, pp. 685–707, Mar. 2005, doi: 10.1007/s00018-004-4513-1.

[88] G. D. Havemann, E. M. Sampson, and T. A. Bobik, ‘PduA is a shell protein of polyhedral organelles involved in coenzyme B(12)-dependent degradation of 1,2-propanediol in Salmonella enterica serovar typhimurium LT2’, J. Bacteriol., vol. 184, no. 5, pp. 1253–1261, Mar. 2002, doi: 10.1128/JB.184.5.1253-1261.2002.

[89] N. W. Kennedy, S. P. Ikonomova, M. Slininger Lee, H. W. Raeder, and D. Tullman-Ercek, ‘Self-assembling Shell Proteins PduA and PduJ have Essential and Redundant Roles in Bacterial Microcompartment Assembly’, J. Mol. Biol., vol. 433, no. 2, p. 166721, Jan. 2021, doi: 10.1016/j.jmb.2020.11.020.

[90] E. M. Sampson and T. A. Bobik, ‘Microcompartments for B12-dependent 1,2-propanediol degradation provide protection from DNA and cellular damage by a reactive metabolic intermediate’, J. Bacteriol., vol. 190, no. 8, pp. 2966–2971, Apr. 2008, doi: 10.1128/JB.01925-07.

[91] E. E. Hansen et al., ‘Pan-genome of the dominant human gut-associated archaeon, Methanobrevibacter smithii, studied in twins’, Proc. Natl. Acad. Sci. U. S. A., vol. 108 Suppl 1, pp. 4599–4606, Mar. 2011, doi: 10.1073/pnas.1000071108.

[92] C. M. Chibani et al., ‘A catalogue of 1,167 genomes from the human gut archaeome’, Nat. Microbiol., vol. 7, no. 1, pp. 48–61, 2022, doi: 10.1038/s41564-021-01020-9.

[93] H. Acar Kirit, J. P. Bollback, and M. Lagator, ‘The Role of the Environment in Horizontal Gene Transfer’, Mol. Biol. Evol., vol. 39, no. 11, p. msac220, Nov. 2022, doi: 10.1093/molbev/msac220.

[94] D. J. Rigden and M. Y. Galperin, ‘Sequence analysis of GerM and SpoVS, uncharacterized bacterial “sporulation” proteins with widespread phylogenetic distribution’, Bioinformatics, vol. 24, no. 16, pp. 1793–1797, Aug. 2008, doi: 10.1093/bioinformatics/btn314.

[95] R. U. Onyenwoke, J. A. Brill, K. Farahi, and J. Wiegel, ‘Sporulation genes in members of the low G+C Gram-type-positive phylogenetic branch (Firmicutes)’, Arch. Microbiol., vol. 182, no. 2, pp. 182–192, Oct. 2004, doi: 10.1007/s00203-004-0696-y.

[96] ‘BacMap’. http://bacmap.wishartlab.com/organisms/525 (accessed Jan. 25, 2023).

[97] S. Nelson-Sathi et al., ‘Origins of major archaeal clades correspond to gene acquisitions from bacteria’, Nature, vol. 517, no. 7532, pp. 77–80, Jan. 2015, doi: 10.1038/nature13805.

[98] A. Ruaud et al., ‘Syntrophy via Interspecies H2 Transfer between Christensenella and Methanobrevibacter Underlies Their Global Cooccurrence in the Human Gut’, mBio, vol. 11, no. 1, pp. e03235-19, Feb. 2020, doi: 10.1128/mBio.03235-19.

[99] S. Varrette, P. Bouvry, H. Cartiaux, and F. Georgatos, ‘Management of an academic HPC cluster: The UL experience’, in 2014 International Conference on High Performance Computing Simulation (HPCS), Jul. 2014, pp. 959–967. doi: 10.1109/HPCSim.2014.6903792.

[100] F. Meyer et al., ‘The metagenomics RAST server – a public resource for the automatic phylogenetic and functional analysis of metagenomes’, BMC Bioinformatics, vol. 9, no. 1, p. 386, Sep. 2008, doi: 10.1186/1471-2105-9-386.

