## Supplementary materials for "Functional prediction of proteins from the human gut archaeome"

### Functional prediction of select proteins from the gut archaeomes - Supplementary materials

### Enhancing annotations of proteins encoded by *Methanobrevibacter smithii*

#### Homologous archaeal proteins

##### Sourmash cluster h1

Annotated as 4-amino-4-deoxy-L-arabinose transferase or related glycosyltransferases of PMT family

4-amino-4-deoxy-L-arabinose transferase (ArnT) is an inner membrane protein that catalyses the covalent addition of the cationic sugar 4-amino-4-deoxy-L-arabinose (L-Ara4N) groups to lipid A, the lipid component of bacterial lipopolysaccharide residing on outer membranes of some bacteria [1], [2]. Glycosylation of the lipid A with L-Ara4N is a common defence strategy described in bacteria, which helps them to resist the action of antimicrobial peptides and polymyxin-class antibiotics [3].

##### Sourmash cluster h2

Annotated as Dolichyl-phosphate-mannose – protein mannosyltransferase 1

Dolichyl-phosphate-mannose–protein mannosyltransferase 1 (POMT1) is involved in protein O-glycosylation, an essential posttranslational modification in eukaryota and certain bacteria [4]. POMT1 transfers mannose residues from dolichyl phosphate-D-mannose to serine/threonine residues of other proteins. All POMTs are integral membrane proteins and form protein complexes which are crucial for mannosyltransferase activity. POMTs are suggested to be evolutionarily related to oligosaccharyltransferases (OSTs) [5]. However, according to literature, not O-linked but N-linked glycosylation occurs in archaea. Some studies report that N-glycosylation is not essential for archaeal cell survival, however, it possibly provides adaptation to severe environments via maintaining S-layer stability and a number of flagella associated with higher motility [6].

##### Sourmash cluster h3

Annotated as Dolichyl-phosphate-mannose – protein mannosyltransferase 1

See sourmash cluster h2

##### Sourmash cluster h4

Annotated as Dolichyl-phosphate-mannose – protein mannosyltransferase 1

See sourmash cluster h2

##### Sourmash cluster h5

Annotated as Dolichyl-diphosphooligosaccharide – protein glycosyltransferase subunit STT3B

STT3B (STT3 oligosaccharyltransferase complex catalytic subunit B) is a catalytic subunit of human oligosaccharyl transferase (OST) complexes which are responsible for the first step on N-linked glycosylation - the transfer of a defined glycan from the lipid carrier. A yeast paralog of STT3B, Stt3, was shown to be structurally and functionally similar to the eubacterial oligosaccharyltransferase PglB and archaebacterial oligosaccharyltransferase AglB [7], [8]. Assuming that the protein in cluster 7 was identified in an archaeon, we suggest that it is a protein more reminiscent of the archaebacterial oligosaccharyltransferase AglB. The above-mentioned enzyme plays a role in the mechanism of N-linked glycosylation, catalysing the final step of the transfer of the glycan tree to the nascent protein [9]. Moreover, Particularly noteworthy is the fact that AglB transfers polysaccharides from dolichol phosphate rather than dolichol pyrophosphate carriers [10], [11].

#### Sourmash cluster h6

Annotated as Dolichyl-phosphate-mannose--protein mannosyltransferase 1

See sourmash cluster

#### Sourmash cluster h7

Annotated as Propanediol utilisation protein pduA

The propanediol utilization (PDU) metabolosome comprises 22 different types of proteins encoded by genes from the pdu operon. PDU metabolosomes are involved in 1,2-propanediol catabolism, which in the first step produces the toxic and reactive intermediate propionaldehyde, which is further converted to propionyl-CoA and 1-propanol. Therefore, the important contribution of MCs involved in 1,2-propanediol degradation is the protection from DNA and cellular damage by a reactive metabolic intermediate propionaldehyde. In the human intestine 1,2-propanediol is excreted by some saccharolytic bacteria when anaerobically grown on monosaccharides, such as fucose and rhamnose. Additionally, 1,2-propanediol is a critical carbon source for certain pathogens and facilitates their proliferation during intestine inflammation [12], [13]. Known bacteria protect themselves via encapsulating the conversion of 1,2-propanediol to propionaldehyde in self-assembling MCs, thereby preventing cellular toxicity and carbon loss [14].

#### Sourmash cluster h9

Annotated as Stage V sporulation protein AE

Sporulation is one of the stages of the bacterial life cycle, when spores are produced in response to the stress of poor nutritional conditions. The spoV loci are involved in the assembly of the spore coat proteins in the forespore. The spoVA operon contains five ORFs, and their functions are not characterised to date [15], [16]. Stage V sporulation protein AD (SpoVAD) is membrane-associated [17] and is essential for dipicolinic acid (DPA) movement across the inner membrane [18]. SpoVAD in *B. subtilis* is predicted to form a channel with other proteins of the spoVA operon through which the major spore component DPA enters the spore in sporulation and exits early in germination. No characterised archaeal species are known for creating endospores. However, sporulation genes are found in a number of non-sporulating bacteria where their function remains unknown too [19].

#### Sourmash cluster h11

Annotated as Phosphoenolpyruvate-dependent PTS system, IIA component

Sugar phosphotransferase system (PTS) is a multi-protein system involved in the regulation of a variety of metabolic and transcriptional processes and is a major carbohydrate transport system in bacteria. Enzyme 2 (EII) is a member of the PTS system and is sugar specific. EII consists of at least three separate subunits IIA, IIB and IIC. archaea do not have a PTS system according to a study from 2002 [20], however in 2012 and 2014 they were reported to exist in archaea too [21], [22].

#### Sourmash cluster h12

Annotated as Immunity protein 7

Immunity protein 7 is a colicin immunity protein that prevents DNase activity of colicin via binding which blocks major dsDNA binding sites. Colicins are bacterial secreted antimicrobial proteins that are produced by E. Coli and kill related bacterial species to reduce competition from them.

#### Sourmash cluster h14

Annotated as Glycerol-3-phosphate cytidylyltransferase

All known glycerol cytidylyltransferase proteins (GCTs) are of bacterial origin and are key to the synthesis of one of the components of teichoic acid. Teichoic acid is a glycopolymer and its main function is to provide flexibility and structural integrity of the bacterial cell wall.

However, genes coding for putative GCTs were found in 2 archaeal organisms *A. fulgidus* and *Aq. aeolicus* [23]. It was shown that discovered GCTs possess stereospecific properties and have preferences for different enantiomers of glycerol phosphate. And different enantiomers of glycerol phosphate in the membrane phospholipids of bacteria and archaea is the most distinctive feature of the two domains [24].

#### Sourmash cluster h15

Annotated as 78 kDa glucose-regulated protein

GRP78 has multiple functions in maintaining cell viability. GRP78 can be located at the plasma membrane where it is cytoprotective [25]. The structure reported by AF2-based ProFunc prediction is matched with the GRP78 ATPase domain. GRP78 consists of two major functional domains: the ATPase (GRP78ATPase) and protein/peptide-binding domains. The ATPase domain has potential as a transmembrane delivery system of cytotoxic agents [26].

#### Sourmash cluster h16

Annotated as Protein from LytR-Cps2A-Psr (LCP) protein family

Domain analysis shows that protein belongs to LytR-Cps2A-Psr (LCP) protein family. LytR-CpsA-Psr family proteins are putative transmembrane proteins present in all Gram-positive organisms, except for the cell wall-deficient *Mollicutes* and one *Clostridiales* strain [27]. LCP proteins were shown to be involved in cell envelope maintenance and bacterial virulence [28], [29]. However, the exact role of LytR-Cps2A-Psr proteins is unknown.

#### Sourmash cluster h18

Annotated as Protein from LytR-Cps2A-Psr (LCP) protein family

See sourmash cluster h16

#### Sourmash cluster h19

Annotated as Uronate isomerase (BH0493)

Uronate isomerase is one of the most divergent members of the amidohydrolase superfamily, which catalyses the reversible isomerization of D-glucuronate to D-fructuronate and D-galacturonate to D-tagaturonate [30]. Uronate isomerase BH0493 is a distinct protein with <20% sequence similarity to any other characterised uronate isomerases from another species than Bacillus halodurans [31]. It was shown that BH0493 can use both D-glucuronate and D-galacturonate as substrates with a similar efficiency.

Glucuronate is a sugar acid derived from glucose. It serves as a building block of glycoglycerolipids and proteoglycans. Glycoglycerolipids are especially abundant in archaea [32]. Galacturonate is also the main component of pectin, an abundant compound in plant biomass. Despite known galacturonate catabolism pathways in bacteria, there has been no proof of such pathways in archaea.

#### Sourmash cluster h20

Annotated as Stage V sporulation protein AD

See cluster h9

#### Sourmash cluster h21

Annotated as Site-specific integrase

Archaeal integrases catalyse reactions beyond site-specific recombination, but also low-sequence specificity recombination which generates deep rearrangements of the host genome [33].

#### Sourmash cluster h25

Annotated as SCO1 protein homolog

SCO1(cytochrome c oxidase assembly protein) is a eukaryotic protein. It is a copper metallochaperone that is located in the inner mitochondrial membrane and is important for the maturation and stabilisation of cytochrome c oxidase subunit II, which plays a role in the regulation of copper homeostasis and transport. According to some studies [20], the Sco1 protein represents the first structure of the class of proteins present in a variety of eukaryotic and bacterial organisms, and elucidates a link between copper trafficking proteins and thioredoxins. Studies suggest that desulfoferrodoxin behaves as a superoxide reductase enzyme and thus provide new insights into the biological mechanisms designed for protection from oxidative stresses [34].

#### Sourmash cluster h26

Annotated as Transmembrane protein 43

The only structure annotation is of probable match likelihood, it has 22.8% identity with the query model. It reports a transmembrane protein 43 (TMEM43), which is a eukaryotic protein with no known homologs in prokaryotes.

#### Sourmash cluster h27

Annotated as GDP-mannose-3', 5’-epimerase

In this case we cannot straightforwardly conclude on the actual function of the protein from provided annotations following our criteria described in *Annotation Selection* (see *Materials and Methods*). In this case both TR and AF predict models with *very high* likelihood and *certain match* significance score, respectively, however, they report different protein names and corresponding structural matches. Therefore, out of four top-mathiching templates (see *Supplementary*) we select the structural model with the highest Identity (25.4% against 25%, 21.99%) and Coverage (98.6% against 96.6%, 76.3%), which in turn corresponds to the 2C5A PDB model GDP-mannose-3', 5’-epimerase.

GDP-mannose 3,5-epimerase (GM35E) belongs to the short-chain dehydrogenase/reductase protein superfamily and catalyses the conversion of GDP-D-mannose towards GDP-L-galactose (GDP-L-Gal) and GDP-L-gulose (GDP-L-Gul). Therefore, this enzyme takes part in the production of rare L-sugars [35]. In bacteria this enzyme is involved in lipopolysaccharide synthesis and in plants it is a key enzyme in vitamin C biosynthesis [36]. It was also shown that in the thermophilic archaeon *T. Acidophilum* GM35E-like enzyme was involved in gulose biosynthesis similar to plant ascorbate biosynthesis [37].

#### Sourmash cluster h28

Annotated as Transthyretin-like protein

Transthyretin (TTR) is a protein responsible for the distribution of thyroid hormones and identified only in vertebrates, however, homologues of the transthyretin-related protein are found in a wide range of species including nonvertebrate species. For instance, PucM is a protein found in many prokaryotes and having high sequence similarity to functionally unrelated transthyretin [38]. MSA built with these 3 proteins (query, TTR and pucM) and the respective phylogenetic tree demonstrate that the query protein does not have any convincing sequence similarity to the other proteins and is placed remotely to both of them on the tree. However, the predicted structure of the query protein was reported to have substantial structure-similarity to TTR. Indeed, tertiary, and quaternary features of the three-dimensional structure of TTR and TTR-like proteins are most likely preserved according to Eneqvist et al [39]. It agrees with the finding that TTR homologs of no obvious function, called TTR-related proteins, have also been identified among various eukaryotic and prokaryotic taxa and constitute a very old and conserved protein family that existed before the divergence of eukaryotes and prokaryotes [39], [40]. TTR and TTR-related proteins have uricase activity and therefore participate in urate catabolism.

#### Sourmash cluster h29

Annotated as Dicer-like 3 protein

Dicer-like 3 protein belongs to the ribonuclease III protein family. It is involved in the generation of small RNAs. Prokaryotic RNAse III is important in post-transcriptional control of mRNA stability and translational efficiency. It is involved in the processing of ribosomal RNA precursors. Prokaryotic RNAse III also plays a role in the maturation of tRNA precursors and in the processing of phage and plasmid transcripts [41].

#### Sourmash cluster h30

Annotated as Interleukin enhancer-binding factor 3

Interleukin enhancer-binding factor 3 (ILF3) is a eukaryotic dsRNA-binding protein, which has no described homologs in prokaryotic organisms. Therefore, based on the results we have for this protein, we can conclude it is a dsRNA-binding protein. Due to the same KEGG annotation for sourmash clusters 41 and 42, one can conclude it contains the same domain, therefore proteins of these two sourmash clusters possess a certain degree of sequence-similarity. Moreover, both proteins are annotated as a Ribonuclease III protein in addition to other template matches. Therefore, the protein of sourmash cluster 42 might have RNase III activity.

#### Sourmash cluster h31

Annotated as 2-aminoethylphosphonate – pyruvate aminotransferase

2-aminoethylphosphonate (2-AEP) – pyruvate aminotransferase is involved in phosphonate degradation and catalyses the reaction of 2-aminoethylphosphonate and pyruvate to 2-phosphonoacetaldehyde and L-alanine. In some bacteria this reaction is the first step in the AEP degradation pathway. 2-phosphonoacetaldehyde is further degraded to phosphonate, which is used by bacteria as a bioavailable source of carbon, nitrogen and phosphorus [42], [43]. Phosphonate has been shown to be produced in all three domains of life, however only in one archaeal species *N. maritimus*. Moreover, *N. maritimus* produces not 2-AEP but 2-hydroxyethylphosphonate (2-HEP), a common intermediate in phosphonate biosynthetic pathways.

#### Sourmash cluster h32

Annotated as PhoP response regulator

All 3D-based results point at a response regulator, and the TR template hit with the highest coverage and identity suggests a PhoP response regulator. Two-component regulator systems (TCRSs) exist in archaea and bacteria. TCRSs are important mediators of signal transduction that enable organisms to detect physical and chemical changes and then relay this signal to modulate gene expression.

PhoP belongs to the OmpR/PhoB family of RRs, which is the largest family and contains thousands of proteins. Despite extensive research in recent years, the molecular mechanism of DNA sequence recognition by this large family of RRs is not fully understood. Euryarchaeota organisms (e.g., halophiles, methanogens, thermophiles) are described to, like bacteria and Eukaryotes, rely on bacterial-type two-component signal transduction systems [44]. PhoP is particularly involved in adaptations to low $Mg^{2+}$ environments and the control of acid resistance genes [45]. However, archaeal TCRSs are different from the bacterial ones, and were hypothetically acquired by archaea from bacteria via HGT [46].

#### Sourmash cluster h35

Annotated as CRISPR system endoribonuclease Csm6

CRISPR-Cas systems provide bacteria and most archaea with defence mechanisms against exogenic genetic elements. Csm6 is a ribonuclease and is associated with type III-A CRISPR–Cas system. Type III systems are thought to be the most ancient type of CRISPR-Cas systems [47]. Csm6 proteins were shown to act as metal-independent endoribonucleases relying on the RNase active site of the HEPN domain for cleavage of ssRNA molecules [48].

#### Sourmash cluster h38

Annotated as Recombination protein uvsY

Homologous recombination (HR) is an important mechanism to provide genome stability and repair and is used for high-precision repair of DNA double-strand breaks. The crucial homology search and DNA strand exchange steps of HR are catalysed by presynaptic filaments - helical filaments of a recombinase enzyme bound to single-stranded DNA [49], [50]. UvsY is required in proper formation of the presynaptic filament.

#### Sourmash cluster h39

Annotated as Intimin/invasin-like protein

Intimins/invasins are members of a broad family of virulence-related bacterial adhesins, sharing a common structure which includes N-terminal β-barrel domain and a C-terminal surface localised passenger domain. Intimins and invasins are outer membrane proteins which interact with the outer environment and promote bacterial/archaeal attachment or invasion [51]. Intimins and invasins are often described in Gram-negative bacteria, however anchorless surface-located adhesins and invasins have been reported in Gram-positive organisms and represent a new class of virulence factors [52].

#### Sourmash cluster h40

Annotated as PTS system, IIB component protein

See sourmash cluster 25

#### Sourmash cluster h41

Annotated as Collagen alpha-1(x) chain (q03692)

Collagen alpha-1(X) chain is a short chain human protein. In bacteria, collagen-like (CL) molecules are cell surface molecules and are found in several pathogenic and non-pathogenic species and are thought to help an alien organism to evade the host immune system [53]–[55]. Studies of archaeal organisms do not include characterised CL proteins.

#### Sourmash cluster h43

Annotated as Hypothetical EmrR homolog

EmrR (E. coli multidrug resistance regulator), also known as MprA (microcin production regulation, locus A), is a member of the MarR protein family of transcriptional regulators. EmrR acts as a negative regulator of the expression of the *emrAB* operon known for multidrug resistance and inducible by metals and antibiotics [56], [57].

### Unique archaeal proteins

#### Sourmash cluster u1

Annotated as Type II restriction endonuclease BglII

Type II restriction endonuclease BglII is a component of the restriction-modification system widespread in prokaryotes. Its role includes protection of the host genome against invading foreign DNA. Although type II restriction enzymes are known to occur in bacterial and archaeal genomes, in the literature there's no information about BglII in archaea [58].

#### Sourmash cluster u2

Annotated as Intimin/invasin-like protein with a Ig-like domain

See sourmash cluster h39

Ig-like domain is widely distributed and present in eukaryotes, plants, fungi, parasites, bacteria, and viruses [59]. Ig-like domains are often found in fimbrial organelles and various cell surface proteins including intimin/invasin family of outer membrane adhesins [60]. Thus, it might play a role in host cell adhesion and invasion of pathogenic strains.

#### Sourmash cluster u3

Annotated as Intimin/invasin-like protein

See sourmash cluster h39, u2

#### Sourmash cluster u4

Annotated as *Unr protein*

AF-based annotation suggests a probable match with a Unr protein, a cytoplasmic RNA-binding protein participating in messenger RNA regulation and stability. The Unr protein contains five cold-shock domains, which are involved in transcriptional and post-transcriptional control of gene expression. Regulated gene expression is achieved via induced response to stress such as nutritional starvation [61].

#### Sourmash cluster u7

Annotated as VIT1/CCC1 transporter family protein

One group of metal transporter proteins in the vacuolar iron transporter (VIT) family, poorly understood proteins that function in secondary active transport of iron across membranes. Homologous proteins exist in plants, yeast, eukaryotes (but not animals) and in bacteria and archaea [62].

Iron is one of the micronutrients essential for microbial growth, development, and physiology. Such metals as iron, nickel, cobalt and others are essential trace elements for methanogens [63]. Iron is in turn a key component of archaeal proteins responsible for energy metabolism and is omnipresent in methanogenic archaea [64].

#### Sourmash cluster u8

Annotated as PilC, type IVa pilus

Ten highly conserved proteins are known to constitute the type 4 pilus (T4P) system. PilC is a motor subunit of this machine [65]. T4P is a type of adhesion filament and exists in nearly all phyla in both prokaryotic domains [66]. T4P carry out different roles, namely cell adhesion, biofilm formation, DNA uptake, protein secretion, exchange of genetic material and species-specific interactions [67]. This mechanism allows archaea rapid adaptation to changing environments. Gene synteny reveals presence of an archaeal flagellar protein FlaI (K07332), homologous to bacterial type IV pili ATPase [68], downstream to u8.

#### Sourmash cluster u12

Annotated as Class 2 OLD family nuclease

Overcoming lysogenization defect (OLD) proteins are a group of uncharacterized nucleases occurring in bacteria, archaea, and some viral genomes. However, speculations suggest that these proteins can be involved in DNA repair and/or replication [69].

#### Sourmash cluster u14

Annotated as Succinate-semialdehyde dehydrogenase

Succinate-semialdehyde dehydrogenase (SSADH) is an enzyme from the aldehyde dehydrogenase protein family and participates in glutamate and butyrate metabolism [95]. SSADH is found in organisms ranging across the tree of life from bacteria to Eukaryotes. An enzyme with succinate semialdehyde activity was characterised in a thermoacidophilic archaeon *S. tokodaii* and was suggested to practically function as succinate semialdehyde dehydrogenase [70]. L-Glutamate can then serve in the colon lumen as a precursor for butyrate and acetate in bacteria. L-Glutamate, in addition to fibre and digestion-resistant starch, can thus serve as a luminally derived fuel precursor for colonocytes [71].

#### Sourmash cluster u15

Annotated as Rubrerythrin

Rubrerythrin is a protein with inorganic pyrophosphatase activity and a C-terminal domain homologous to rubredoxin [72]. FdhE, formate dehydrogenase formation protein, is sought to be an iron-binding rubredoxin and interacts with the catalytic molybdoprotein subunits of both respiratory formate dehydrogenases N and O [73]. Rubrerythrins take part in oxidative stress defence as peroxide scavengers in a wide range of organisms [74].

#### Sourmash cluster u21

Annotated as Pyruvyl transferase 1

Despite pyruvyl transferase is reported by TR template hits with the lowest e-value and the highest z-score, and via the BLAST search based on AF prediction, one may still be not sure, especially due to the AF-based structural hit to a functionally uncharacterised protein. Other TR templates suggest matches with other proteins with transferase activity (i.e., glycogen phosphorylase, glycosyltransferase family 9).

Pyruvyl-transferases are involved in peptidoglycan-associated polymer biosynthesis [75]. Although archaea lack peptidoglycan (or murein) like Eukaryotes, methanogens instead use pseudomurein, structurally similar to the peptidoglycan. Besides that, archaeal cell wall polymers include methanochondroitin, named for its similarities to vertebrate chondroitin. The unique structure of archaeal cell walls confers an advantage in resistance against high internal osmotic pressure [76].

#### Sourmash cluster u22

Annotated as Type I restriction-modification EcoKI enzyme, specificity subunit

Restriction–modification (R–M) systems are protein complexes which protect the host bacterium from invasion by foreign DNA through global methylation by methyltransferase (MTase) activity and digestion of the invaded DNA by restriction endonuclease (REase) activity [77], [78]. Amongst the R–M systems, the Type I R–M systems in particular have been shown to be important for gene regulation and possibly pathogen virulence.

A complete type I R-M system consists of a specificity (S), a modification (M) and a restriction (R) subunit. Subunits encoded M and S are both necessary and sufficient for methyltransferase activity [78]. In bacteria the type I R-M systems have been shown to provide a barrier to phage infection in the host, allow for further phage defence and possibly for regulatory control [79], [80]. The EcoKI type I R-M systems modulate the exchange of genetic material between microbial cells and plasmids, viruses, and transposons. R-M system EcoKI was shown to play significant role in acceptance or rejection of genetic material transferred by conjugation of microbial cells [81].

#### Sourmash cluster u23

Annotated as Sulfur carrier protein TtuB

This protein is involved in thiolation, a widely conserved in all domains of life post transcriptional modification. One of such modifications, 2-thiouridine (s2U), enhances thermostability in thermophilic bacteria. One of the proteins involved in this process, 2-thiouridine synthesis sulfur carrier protein TtuB, functions as a sulfur donor in the sulfurtransferase reaction [82]. However, up-to-date this protein is only described in bacteria, while in archaea another protein functioning as a sulfur carrier for 2-thiouridine synthesis and a protein modifier, SAMP2, is known [83].

#### Sourmash cluster u24

Annotated as Polypeptide N-acetylgalactosaminyltransferase

Polypeptide N-acetylgalactosaminyltransferase (GalNAc) in a member of a large family of glycosyltransferases [84]. This enzyme is involved in O- and N-glycosylation, in particular in the UDP-GlcNAc biosynthetic pathway. In archaea, the bacterial-type UDP-GlcNAc biosynthetic pathway was reported for *M.* maripaludis [85].

#### Sourmash cluster u25

Annotated as Nickel (III) ABC transporter, periplasmic iron-binding protein

Periplasmic binding proteins exist in Gram-negative bacteria and are dissolved in the periplasm, while in archaea and Gram-positive bacteria such proteins are external membrane-bound lipoproteins [86], [87]. These proteins participate in recognition processes and provide chemoreception and transmembrane transport. The transport of solute molecules into the cytoplasm is ensured via ABC transporters. Nickel is one of the essential trace elements used by prokaryotes, and performs key functions in various metabolic processes, such as energy metabolism and virulence. Nickel was shown to be used by the methanogenic archaeon *M. thermoautotrophicum* for growth [88].

#### Sourmash cluster u27

Annotated as ARP1 actin related protein

Actin-related proteins (Arps) are a class of proteins found in all eukaryotes and many species of bacteria and archaea. Conventional actin assembles into two-stranded helical filaments (which reminds the shape predicted for this archaeal protein) that form structural scaffolds used to organise the intracellular space and to drive cell division, shape change, and cell locomotion. Prokaryotic Arps were discovered only recently, and their functions are not well understood. Arp1 is a cytoskeleton-associated Arp and regulates the assembly and function of the actin and microtubule cytoskeletons [89]. Arp1 forms a short filament as part of the dynactin complex that promotes cargo movement by the microtubule motor dynein [90]. Golgin-mediated tethering is thought to be important for vesicular traffic at the Golgi apparatus, the maintenance of Golgi architecture, as well as the positioning of the Golgi apparatus within cells. Although it is now established that golgins are membrane and cytoskeleton tethers, the mechanisms underlying tethering remain poorly defined [91].

#### Characterization of select proteins and gene structures in *Methanobrevibacter smithii* genomes

We describe functions of the available proteins and distinguish the most frequently occurring processes. Our results demonstrate that homologous proteins from clusters *h1, h2, h3, h4, h5* and *h6* constitute the most prevalent archaeal gut-specific process and thus deserve particular attention. All proteins of this clusters are types of glycosyltransferases (GTs) acting on various sugar donors via transferring sugar residues from donors to acceptor molecules through establishing natural glycosidic linkages. GTs are also described to be the most common enzyme class in archaea [92], yet hyperthermophilic archaeal species encode fewer glycosyltransferases than nonhyperthermophilic archaea [93]. Gene synteny analysis reveals that the flanking regions of the abovementioned genes also contain various transferases. In our work, the most prevalent unique archaeal protein is from the group of GTs from cluster *h1* and is a *4-amino-4-deoxy-L-arabinose (L-Ara4N) transferase*. The mentioned protein was predicted as L-Ara4N transferase with both tools (AF+PF and TR) with *very high* confidence score, therefore we can assume that the homology behind this annotation is robust. This protein is reported to transfer a L-Ara4N unit to the lipid A, a complex glycolipid on the surface of most Gram-negative bacteria, which is essential for protection from environmental stress, symbiosis, virulence and resistance against antimicrobial activity [94], [95]. Whether it encodes for similar functions in archaea, given their unique cell wall composition [96], which is unlike Gram-negative bacteria, needs to be validated in future studies. Moreover, one of the six GTs is a *dolichyl-diphosphooligosaccharide-protein glycosyltransferase subunit STT3B* (*h5*) and functions as an accessory protein in N-glycosylation and provides its maximal efficiency [97]. Indeed, N-linked glycosylation is known as a ubiquitous posttranslational modification in archaea, however this fact has only been confirmed in a limited number of species. For example, N-linked glycans have been observed in methanogens such as *M. voltae [98], M. maripaludis* [99], *M. fervidus* [100]. Although there is yet no evidence about N-linked glycosylation pathways in *M. smithii* or other *Methanobrevibacter* species, multiple related studies point to its probable existence. Indeed, archaeal N-glycosylation plays an important role for the viability and adaptivity of archaeal cells to external conditions such as high salinity [101], [102], elevated temperatures [100] acidic environment [103], it also maintains structural integrity of cells [104], [105]. Similarly, N-glycosylation of archaeal proteins might benefit species' viability and adaptation to the intestinal conditions.

Four out of six identified GTs (clusters *h2, h3, h4, h6*) are *dolichyl-phosphate-mannose-protein mannosyltransferases 1 (POMT1),* which are responsible for O-linked glycosylation of proteins in eukaryotes. However, O-glycosylation occurring in archaea is not well understood. There is limited evidence about O-glycosylation in archaea, yet it is known that threonine-rich regions of S-layer glycoproteins supplement galactose–glucose disaccharides in *Hbt. salinarum* and *Hfx. Volcanii* [106], [107]. However, another O-glycosylation associated protein, *polypeptide N-acetylgalactosaminyltransferase*, was found in the subset of unique archaeal proteins (unique cluster u24). O-glycosylation is mainly found in eukaryotic cells. In particular, mucins are examples of O-glycosylated glycoproteins. Mucins are the main structural components of the intestinal mucus and contribute to the interaction between gut microorganisms and epithelial surfaces [108]. On the other hand, Samuel *et al.* demonstrated that *M. smithii* decorates its cellular surface with sugar residues mimicking those present in the glycan landscape of the intestinal environment [109]. Taken together, the fact that *M. smithii* has human mucus- and epithelial cell surface-associated glycans and that archaeal gut species possess coding potential for enzymes which function in O-linked glycosylation suggests that *M. smithii* cells could indeed mimic surfaces of eukaryotic cells of the intestinal mucus. O-glycans do not only play a role on the structural level of individual proteins, but also O-glycans might modulate protein interactions between other proteins as well as cells, which in turn benefits cell trafficking and environmental recognition [110]. O-glycosylated proteins are traditionally known to be localized extracellularly including cell surfaces and luminal compartments [111]. However, Torres and Hart demonstrated that *O*-linked modification is dynamic, reversible and short lived and O-linked GlcNAc residues often occur intracellularly [112]. Therefore, O-glycosylated proteins might also have regulatory functions. With regards to the former statement, Van den Steen *et al.* report that two main proteins implicated in Alzheimer's disease – tau and beta-amyloid precursor protein – are known to be O-glycosylated and increased in disease [113].

Besides POMT1 and related GTs, there are other archaeal proteins which reflect adaptation to external conditions and cell survival (clusters *h7, h12, h15, h21, h25, h35* and *h38*). Interestingly, all these proteins are considerably enriched in archaea rather than bacteria based on their relative occurrence. For example, the second most frequently occurring archaeal protein (h7) in the group of homologous proteins is structurally similar to *propanediol utilization protein pduA. PduA* has yet been described only in bacteria. *PduA* is a major shell protein that is essential for bacterial microcompartment assembly and required for the formation of polyhedral organelles involved in coenzyme B12-dependent 1,2-propanediol degradation. The majority of bacterial microcompartments (MCs) are metabolosomes. It is hypothesized that the primary role of MCs involved in 1,2-propanediol degradation might be the regulation of aldehyde production to prevent toxicity and DNA damage [114]–[116]. Metabolosomes are also found in archaea including those inhabiting human intestine [117]. Even though 1,2-propanediol degrading MCs have not been described in archaea up to date, detection of proteins involved in such a process suggests that gut archaea might somehow protect themselves from toxicity exhibited by surrounding organisms and even pathogens. For instance, in the human intestine 1,2-propanediol is excreted by some saccharolytic bacteria when anaerobically grown on monosaccharides, such as fucose and rhamnose [12], [13].

Protein clusters that are relatively equally represented by archaeal and bacterial proteins (clusters *h11*, *h27* and *h31*) are all involved in energy metabolism and related processes. Indeed, archaeal metabolism resembles in its complexity the metabolism of bacteria and lower Eukaryotes [118]. For instance, cluster *h31* is represented by *2-aminoethylphosphonate-pyruvate (2-AEP) aminotransferase*, a very common bacterial enzyme playing a role in phosphonate degradation [119]. Phosphonate degradation has also been shown as an important source and a common production pathway for methane [120]. More specifically, Sosa *et al.* demonstrated that phosphonate degradation is a source and production pathway of methane and ethylene in surface waters of western North Atlantic Ocean [121]. We hypothesize *M. smithii* might degrade 2-AEP produced by bacteria [122], pointing at symbiotic relationships between archaea and bacteria, however the role of 2-AEP aminotransferase in archaea is yet not described in the literature [123]. In general, the exact biological function of phosphonate macromolecules has not been comprehensively described [124]. Cluster *h11* represents a *phosphoenolpyruvate-dependent sugar phosphotransferase system (PTS) system, IIA component*. Until recently, it was assumed that PTS proteins are absent from archaeal species except for their presence in *H. marismortui* [125]. However, today it is widely accepted that archaea possess PTS, and it is an essential player for key cellular processes. In addition, several proteins performing metabolic functions are more specific to archaea (homologous clusters h19, h28), for example, the protein from cluster *h28* is a *transthyretin-like protein*. It has been shown that modern transthyretins evolved from their ancient ancestors with a functional role in uric acid catabolism through an event of duplication followed by divergence [126], [127]. By assuming that the described evolutionary history of transthyretin (TTR) and TTR-related proteins is indeed correct, we speculate that the archaeal protein identified in our work might have uricase activity and therefore participate in urate catabolism. Functional uricase enzymes are found in eukaryotes and prokaryotes. There exist only limited studies describing uricase activity in archaea, however, it was shown that the strict anaerobe, *M. vannielii*, among certain purines can use uric acid as a sole nitrogen source for growth [128]. According to other studies, uric acid fermentation might be a triggering step for methane production in geese [129].

The prevalence of unique archaeal protein clusters is significantly lower in the gut compared to archaeal-bacterial homologs. However, the most frequent ones also play a role in adaptation and host cell defense processes. The most frequent protein cluster is the *type II restriction endonuclease BglII* (*u1*), whose primary function is the protection of the host genome against foreign DNA, for example invading viral DNA. Another example of such a defense mechanism is demonstrated by the presence of the *specificity unit of type I restriction-modification (R-M) EcoKI enzyme* (*u22*). Roer *et al.* report that *R-M EcoKI enzyme* is involved in regulation of rejection/acceptance of genetic material transferred upon conjugation of microbial cells [81]. Conjugation is known as a mechanism of horizontal gene transfer which plays an important role in the evolution of archaeal species and provides means of adaptation to new ecological niches [130]–[132]. Indeed, archaea are known to encode genes of bacterial origin, moreover the flow of genes from bacteria to archaea occurs several-fold more often than in the reverse direction [133].

Unique cluster *u4* is represented by the *Unr protein* involved in stress adaptation. Cold-shock domains comprising this protein provide adaptation to external conditions through stress-induced transcriptional and post-transcriptional control of gene expression. Several frequently occurring archaeal proteins were annotated as adhesin-like, and such proteins were found in both bacterial homologous and unique groups, however the unique archaeal counterparts are more frequent in the gut (*h39*, *u2, u3).* It was shown that *M. smithii* possess a variety of adhesin-like proteins used in different repertoires with different substrates. This strategy allows *M. smithii* strains developing adaptive responses to different metabolic niches in the gut, as well as establishing syntrophic relationships with bacterial neighbours with other metabolic potentials [134]. Taken together, these mechanisms contribute to the viability of archaea in the human gut.

### References

[1] V. I. Petrou *et al.*, ‘Structures of aminoarabinose transferase ArnT suggest a molecular basis for lipid A glycosylation’, *Science*, vol. 351, no. 6273, pp. 608–612, Feb. 2016, doi: 10.1126/science.aad1172.

[2] L. E. Bretscher, M. T. Morrell, A. L. Funk, and C. S. Klug, ‘Purification and characterization of the L-Ara4N transferase protein ArnT from Salmonella typhimurium’, *Protein Expr. Purif.*, vol. 46, no. 1, pp. 33–39, Mar. 2006, doi: 10.1016/j.pep.2005.08.028.

[3] B. D. Needham and M. S. Trent, ‘Fortifying the barrier: the impact of lipid A remodelling on bacterial pathogenesis’, *Nat. Rev. Microbiol.*, vol. 11, no. 7, pp. 467–481, Jul. 2013, doi: 10.1038/nrmicro3047.

[4] S. Strahl-Bolsinger, T. Immervoll, R. Deutzmann, and W. Tanner, ‘PMT1, the gene for a key enzyme of protein O-glycosylation in Saccharomyces cerevisiae.’, *Proc. Natl. Acad. Sci. U. S. A.*, vol. 90, no. 17, pp. 8164–8168, Sep. 1993, Accessed: Feb. 16, 2022. [Online]. Available: https://www.ncbi.nlm.nih.gov/pmc/articles/PMC47309/

[5] L. Bai, A. Kovach, Q. You, A. Kenny, and H. Li, ‘Structure of the eukaryotic protein O-mannosyltransferase Pmt1−Pmt2 complex’, *Nat. Struct. Mol. Biol.*, vol. 26, no. 8, Art. no. 8, Aug. 2019, doi: 10.1038/s41594-019-0262-6.

[6] S. Yurist-Doutsch, B. Chaban, D. J. VanDyke, K. F. Jarrell, and J. Eichler, ‘Sweet to the extreme: protein glycosylation in Archaea’, *Mol. Microbiol.*, vol. 68, no. 5, pp. 1079–1084, 2008, doi: 10.1111/j.1365-2958.2008.06224.x.

[7] S. Shrimal and R. Gilmore, ‘Oligosaccharyltransferase structures provide novel insight into the mechanism of asparagine-linked glycosylation in prokaryotic and eukaryotic cells’, *Glycobiology*, vol. 29, no. 4, pp. 288–297, Oct. 2018, doi: 10.1093/glycob/cwy093.

[8] S. Mohanty, B. P. Chaudhary, and D. Zoetewey, ‘Structural Insight into the Mechanism of N-Linked Glycosylation by Oligosaccharyltransferase’, *Biomolecules*, vol. 10, no. 4, p. E624, Apr. 2020, doi: 10.3390/biom10040624.

[9] B. H. Meyer and S.-V. Albers, ‘AglB, catalyzing the oligosaccharyl transferase step of the archaeal N-glycosylation process, is essential in the thermoacidophilic crenarchaeon Sulfolobus acidocaldarius’, *MicrobiologyOpen*, vol. 3, no. 4, pp. 531–543, Aug. 2014, doi: 10.1002/mbo3.185.

[10] C. Kuntz, J. Sonnenbichler, I. Sonnenbichler, M. Sumper, and R. Zeitler, ‘Isolation and characterization of dolichol-linked oligosaccharides from Haloferax volcanii’, *Glycobiology*, vol. 7, no. 7, pp. 897–904, Oct. 1997, doi: 10.1093/glycob/7.7.897.

[11] J. Eichler, ‘Post-translational modification of the S-layer glycoprotein occurs following translocation across the plasma membrane of the haloarchaeon Haloferax volcanii’, *Eur. J. Biochem.*, vol. 268, no. 15, pp. 4366–4373, Aug. 2001, doi: 10.1046/j.1432-1327.2001.02361.x.

[12] B. Dogan *et al.*, ‘Inflammation-associated adherent-invasive Escherichia coli are enriched in pathways for use of propanediol and iron and M-cell translocation’, *Inflamm. Bowel Dis.*, vol. 20, no. 11, pp. 1919–1932, Nov. 2014, doi: 10.1097/MIB.0000000000000183.

[13] C. Fan, S. Cheng, S. Sinha, and T. A. Bobik, ‘Interactions between the termini of lumen enzymes and shell proteins mediate enzyme encapsulation into bacterial microcompartments’, *Proc. Natl. Acad. Sci. U. S. A.*, vol. 109, no. 37, pp. 14995–15000, Sep. 2012, doi: 10.1073/pnas.1207516109.

[14] S. D. Axen, O. Erbilgin, and C. A. Kerfeld, ‘A Taxonomy of Bacterial Microcompartment Loci Constructed by a Novel Scoring Method’, *PLoS Comput. Biol.*, vol. 10, no. 10, p. e1003898, Oct. 2014, doi: 10.1371/journal.pcbi.1003898.

[15] Roy H. Doi, ‘Sporulation and Germination’, in *Bacillus*, Springer, Boston, MA, 1989. Accessed: Feb. 10, 2022. [Online]. Available: https://doi.org/10.1007/978-1-4899-3502-1_8

[16] R. S. Prajapati and S. M. Cutting, ‘10 - Spores, Sporulation and Germination’, in *Molecular Medical Microbiology*, M. Sussman, Ed., London: Academic Press, 2002, pp. 199–208. doi: 10.1016/B978-012677530-3/50229-4.

[17] Y. Chen, W. K. Ray, R. F. Helm, S. B. Melville, and D. L. Popham, ‘Levels of Germination Proteins in Bacillus subtilis Dormant, Superdormant, and Germinating Spores’, *PLOS ONE*, vol. 9, no. 4, p. e95781, Apr. 2014, doi: 10.1371/journal.pone.0095781.

[18] G. Korza and P. Setlow, ‘Topology and accessibility of germination proteins in the Bacillus subtilis spore inner membrane’, *J. Bacteriol.*, vol. 195, no. 7, pp. 1484–1491, Apr. 2013, doi: 10.1128/JB.02262-12.

[19] D. J. Rigden and M. Y. Galperin, ‘Sequence analysis of GerM and SpoVS, uncharacterized bacterial “sporulation” proteins with widespread phylogenetic distribution’, *Bioinformatics*, vol. 24, no. 16, pp. 1793–1797, Aug. 2008, doi: 10.1093/bioinformatics/btn314.

[20] A. I. Slesarev *et al.*, ‘The complete genome of hyperthermophile Methanopyrus kandleri AV19 and monophyly of archaeal methanogens’, *Proc. Natl. Acad. Sci.*, vol. 99, no. 7, pp. 4644–4649, Apr. 2002, doi: 10.1073/pnas.032671499.

[21] A. Pickl, U. Johnsen, and P. Schönheit, ‘Fructose Degradation in the Haloarchaeon Haloferax volcanii Involves a Bacterial Type Phosphoenolpyruvate-Dependent Phosphotransferase System, Fructose-1-Phosphate Kinase, and Class II Fructose-1,6-Bisphosphate Aldolase’, *J. Bacteriol.*, Apr. 2012, doi: 10.1128/JB.00200-12.

[22] L. Cai *et al.*, ‘Analysis of the Transcriptional Regulator GlpR, Promoter Elements, and Posttranscriptional Processing Involved in Fructose-Induced Activation of the Phosphoenolpyruvate-Dependent Sugar Phosphotransferase System in Haloferax mediterranei’, *Appl. Environ. Microbiol.*, Feb. 2014, Accessed: Feb. 03, 2022. [Online]. Available: https://journals.asm.org/doi/abs/10.1128/AEM.03372-13

[23] M. V. Rodrigues, N. Borges, and H. Santos, ‘Glycerol Phosphate Cytidylyltransferase Stereospecificity Is Key to Understanding the Distinct Stereochemical Compositions of Glycerophosphoinositol in Bacteria and Archaea’, *Appl. Environ. Microbiol.*, Oct. 2016, doi: 10.1128/AEM.02462-16.

[24] Y. Koga, ‘From promiscuity to the lipid divide: on the evolution of distinct membranes in Archaea and Bacteria’, *J. Mol. Evol.*, vol. 78, no. 3–4, pp. 234–242, Apr. 2014, doi: 10.1007/s00239-014-9613-4.

[25] C. Casas, ‘GRP78 at the Centre of the Stage in Cancer and Neuroprotection’, *Front. Neurosci.*, vol. 11, 2017, Accessed: Feb. 07, 2022. [Online]. Available: https://www.frontiersin.org/article/10.3389/fnins.2017.00177

[26] S. J. Hughes, T. Antoshchenko, Y. Chen, H. Lu, J. C. Pizarro, and H.-W. Park, ‘Probing the ATP Site of GRP78 with Nucleotide Triphosphate Analogs’, *PLOS ONE*, vol. 11, no. 5, p. e0154862, May 2016, doi: 10.1371/journal.pone.0154862.

[27] J. Hübscher, L. Lüthy, B. Berger-Bächi, and P. Stutzmann Meier, ‘Phylogenetic distribution and membrane topology of the LytR-CpsA-Psr protein family’, *BMC Genomics*, vol. 9, no. 1, p. 617, Dec. 2008, doi: 10.1186/1471-2164-9-617.

[28] C. Stefanović, F. F. Hager, and C. Schäffer, ‘LytR-CpsA-Psr Glycopolymer Transferases: Essential Bricks in Gram-Positive Bacterial Cell Wall Assembly’, *Int. J. Mol. Sci.*, vol. 22, no. 2, Art. no. 2, Jan. 2021, doi: 10.3390/ijms22020908.

[29] Q. Wang *et al.*, ‘CpsA, a LytR-CpsA-Psr Family Protein in Mycobacterium marinum, Is Required for Cell Wall Integrity and Virulence’, *Infect. Immun.*, vol. 83, no. 7, pp. 2844–2854, Jul. 2015, doi: 10.1128/IAI.03081-14.

[30] T. T. Nguyen *et al.*, ‘The Mechanism of the Reaction Catalyzed by Uronate Isomerase Illustrates How an Isomerase May Have Evolved from a Hydrolase within the Amidohydrolase Superfamily’, *Biochemistry*, vol. 48, no. 37, pp. 8879–8890, Sep. 2009, doi: 10.1021/bi901046x.

[31] T. T. Nguyen *et al.*, ‘At the periphery of the amidohydrolase superfamily: Bh0493 from Bacillus halodurans catalyzes the isomerization of D-galacturonate to D-tagaturonate’, *Biochemistry*, vol. 47, no. 4, pp. 1194–1206, Jan. 2008, doi: 10.1021/bi7017738.

[32] M. Kamekura and M. Kates, ‘Structural diversity of membrane lipids in members of Halobacteriaceae’, in *Bioscience, Biotechnology, and Biochemistry*, 6th ed.1999, pp. 969–972. Accessed: Feb. 14, 2022. [Online]. Available: https://doi.org/10.1271/bbb.63.969

[33] C. Badel, V. Da Cunha, and J. Oberto, ‘Archaeal tyrosine recombinases’, *FEMS Microbiol. Rev.*, vol. 45, no. 4, p. fuab004, Jul. 2021, doi: 10.1093/femsre/fuab004.

[34] M. Lombard, M. Fontecave, D. Touati, and V. Nivière, ‘Reaction of the Desulfoferrodoxin from Desulfoarculus baarsii with Superoxide Anion: EVIDENCE FOR A SUPEROXIDE REDUCTASE ACTIVITY*’, *J. Biol. Chem.*, vol. 275, no. 1, pp. 115–121, Jan. 2000, doi: 10.1074/jbc.275.1.115.

[35] O. Gevaert, S. Van Overtveldt, K. Beerens, and T. Desmet, ‘Characterization of the First Bacterial and Thermostable GDP-Mannose 3,5-Epimerase’, *Int. J. Mol. Sci.*, vol. 20, no. 14, p. 3530, Jul. 2019, doi: 10.3390/ijms20143530.

[36] K. Beerens, O. Gevaert, and T. Desmet, ‘GDP-Mannose 3,5-Epimerase: A View on Structure, Mechanism, and Industrial Potential’, *Front. Mol. Biosci.*, vol. 8, 2022, Accessed: Feb. 11, 2022. [Online]. Available: https://www.frontiersin.org/article/10.3389/fmolb.2021.784142

[37] N. Yamauchi and Y. Nakayama, ‘Biosynthetic mechanism for L-Gulose in main polar lipids of Thermoplasma acidophilum and possible resemblance to plant ascorbic acid biosynthesis’, *Biosci. Biotechnol. Biochem.*, vol. 77, no. 10, pp. 2087–2093, 2013, doi: 10.1271/bbb.130442.

[38] A. C. Schultz, P. Nygaard, and H. H. Saxild, ‘Functional Analysis of 14 Genes That Constitute the Purine Catabolic Pathway in Bacillus subtilis and Evidence for a Novel Regulon Controlled by the PucR Transcription Activator’, *J. Bacteriol.*, Jun. 2001, doi: 10.1128/JB.183.11.3293-3302.2001.

[39] T. Eneqvist, E. Lundberg, L. Nilsson, R. Abagyan, and A. E. Sauer-Eriksson, ‘The transthyretin-related protein family’, *Eur. J. Biochem.*, vol. 270, no. 3, pp. 518–532, 2003, doi: 10.1046/j.1432-1033.2003.03408.x.

[40] P. Prapunpoj, K. Yamauchi, N. Nishiyama, S. J. Richardson, and G. Schreiber, ‘Evolution of structure, ontogeny of gene expression, and function of Xenopus laevis transthyretin’, *Am. J. Physiol.-Regul. Integr. Comp. Physiol.*, vol. 279, no. 6, pp. R2026–R2041, Dec. 2000, doi: 10.1152/ajpregu.2000.279.6.R2026.

[41] A. W. Nicholson, ‘Ribonuclease III mechanisms of double-stranded RNA cleavage’, *WIREs RNA*, vol. 5, no. 1, pp. 31–48, 2014, doi: 10.1002/wrna.1195.

[42] A. M. Cook, C. G. Daughton, and M. Alexander, ‘Phosphonate utilization by bacteria’, *J. Bacteriol.*, Jan. 1978, doi: 10.1128/jb.133.1.85-90.1978.

[43] C. C. H. Chen *et al.*, ‘Degradation Pathway of the Phosphonate Ciliatine: Crystal Structure of 2-Aminoethylphosphonate Transaminase’, *Biochemistry*, vol. 41, no. 44, pp. 13162–13169, Nov. 2002, doi: 10.1021/bi026231v.

[44] D. Esser *et al.*, ‘Protein phosphorylation and its role in archaeal signal transduction’, *FEMS Microbiol. Rev.*, vol. 40, no. 5, pp. 625–647, Sep. 2016, doi: 10.1093/femsre/fuw020.

[45] A. Kato, H. Tanabe, and R. Utsumi, ‘Molecular characterization of the PhoP-PhoQ two-component system in Escherichia coli K-12: identification of extracellular Mg2+-responsive promoters’, *J. Bacteriol.*, vol. 181, no. 17, pp. 5516–5520, Sep. 1999, doi: 10.1128/JB.181.17.5516-5520.1999.

[46] T. Krell, 'Exploring the (Almost) Unknown: Archaeal Two-Component Systems', *J. Bacteriol.*, Jan. 2018, doi: 10.1128/JB.00774-17.

[47] P. Mohanraju, K. S. Makarova, B. Zetsche, F. Zhang, E. V. Koonin, and J. van der Oost, 'Diverse evolutionary roots and mechanistic variations of the CRISPR-Cas systems', *Science*, vol. 353, no. 6299, p. aad5147, Aug. 2016, doi: 10.1126/science.aad5147.

[48] K. Foster, J. Kalter, W. Woodside, R. M. Terns, and M. P. Terns, 'The ribonuclease activity of Csm6 is required for anti-plasmid immunity by Type III-A CRISPR-Cas systems', *RNA Biol.*, vol. 16, no. 4, pp. 449–460, Apr. 2019, doi: 10.1080/15476286.2018.1493334.

[49] K. Pant, L. Shokri, R. L. Karpel, S. W. Morrical, and M. C. Williams, 'Modulation of T4 gene 32 protein DNA binding activity by the recombination mediator protein UvsY', *J. Mol. Biol.*, vol. 380, no. 5, pp. 799–811, Jul. 2008, doi: 10.1016/j.jmb.2008.05.039.

[50] J. Liu, K. T. Ehmsen, W.-D. Heyer, and S. W. Morrical, 'Presynaptic Filament Dynamics in Homologous Recombination and DNA Repair', *Crit. Rev. Biochem. Mol. Biol.*, vol. 46, no. 3, pp. 240–270, Jun. 2011, doi: 10.3109/10409238.2011.576007.

[51] T. M. Adams, A. Wentzel, and H. Kolmar, 'Intimin-Mediated Export of Passenger Proteins Requires Maintenance of a Translocation-Competent Conformation', *J. Bacteriol.*, vol. 187, no. 2, pp. 522–533, Jan. 2005, doi: 10.1128/JB.187.2.522-533.2005.

[52] G. S. Chhatwal, 'Anchorless adhesins and invasins of Gram-positive bacteria: a new class of virulence factors', *Trends Microbiol.*, vol. 10, no. 5, pp. 205–208, May 2002, doi: 10.1016/S0966-842X(02)02351-X.

[53] C. C. Caswell, M. Barczyk, D. R. Keene, E. Lukomska, D. E. Gullberg, and S. Lukomski, 'Identification of the First Prokaryotic Collagen Sequence Motif That Mediates Binding to Human Collagen Receptors, Integrins α2β1 and α11β1', *J. Biol. Chem.*, vol. 283, no. 52, pp. 36168–36175, Dec. 2008, doi: 10.1074/jbc.M806865200.

[54] L. N. Waller *et al.*, 'Identification of a Second Collagen-Like Glycoprotein Produced by Bacillus anthracis and Demonstration of Associated Spore-Specific Sugars', *J. Bacteriol.*, vol. 187, no. 13, pp. 4592–4597, Jul. 2005, doi: 10.1128/JB.187.13.4592-4597.2005.

[55] C. Xu, Z. Yu, M. Inouye, B. Brodsky, and O. Mirochnitchenko, 'Expanding the family of collagen proteins: recombinant bacterial collagens of varying composition form triple-helices of similar stability', *Biomacromolecules*, vol. 11, no. 2, pp. 348–356, Feb. 2010, doi: 10.1021/bm900894b.

[56] H. Zhang, Y. Ma, P. Liu, and X. Li, 'Multidrug resistance operon emrAB contributes for chromate and ampicillin co-resistance in a Staphylococcus strain isolated from refinery polluted river bank', *SpringerPlus*, vol. 5, no. 1, p. 1648, Sep. 2016, doi: 10.1186/s40064-016-3253-7.

[57] K. Nishino, S. Yamasaki, R. Nakashima, M. Zwama, and M. Hayashi-Nishino, 'Function and Inhibitory Mechanisms of Multidrug Efflux Pumps', *Front. Microbiol.*, vol. 12, 2021, Accessed: Feb. 18, 2022. [Online]. Available: https://www.frontiersin.org/article/10.3389/fmicb.2021.737288

[58] A. Pingoud and A. Jeltsch, 'Structure and function of type II restriction endonucleases', *Nucleic Acids Res.*, vol. 29, no. 18, pp. 3705–3727, Sep. 2001, doi: 10.1093/nar/29.18.3705.

[59] D. M. Halaby and J. P. Mornon, 'The immunoglobulin superfamily: an insight on its tissular, species, and functional diversity', *J. Mol. Evol.*, vol. 46, no. 4, pp. 389–400, Apr. 1998, doi: 10.1007/pl00006318.

[60] 'Immunoglobulin domains in Escherichia coli and other enterobacteria: from pathogenesis to applications in antibody technologies | FEMS Microbiology Reviews | Oxford Academic'. https://academic.oup.com/femsre/article/37/2/204/623411

[61] V. Dormoy-Raclet *et al.*, 'Unr, a cytoplasmic RNA-binding protein with cold-shock domains, is involved in control of apoptosis in ES and HuH7 cells', *Oncogene*, vol. 26, no. 18, pp. 2595–2605, Apr. 2007, doi: 10.1038/sj.onc.1210068.

[62] S. Bhubhanil, J. Chamsing, P. Sittipo, P. Chaoprasid, R. Sukchawalit, and S. 2014 Mongkolsuk, 'Roles of Agrobacterium tumefaciens membrane-bound ferritin (MbfA) in iron transport and resistance to iron under acidic conditions', *Microbiology*, vol. 160, no. 5, pp. 863–871, doi: 10.1099/mic.0.076802-0.

[63] P. Scherer, H. Lippert, and G. Wolff, 'Composition of the major elements and trace elements of 10 methanogenic bacteria determined by inductively coupled plasma emission spectrometry', *Biol. Trace Elem. Res.*, vol. 5, no. 3, pp. 149–163, Jun. 1983, doi: 10.1007/BF02916619.

[64] D. C. Johnson, D. R. Dean, A. D. Smith, and M. K. Johnson, 'Structure, function, and formation of biological iron-sulfur clusters', *Annu. Rev. Biochem.*, vol. 74, pp. 247–281, 2005, doi: 10.1146/annurev.biochem.74.082803.133518.

[65] C. L. Giltner, Y. Nguyen, and L. L. Burrows, 'Type IV Pilin Proteins: Versatile Molecular Modules', *Microbiol. Mol. Biol. Rev. MMBR*, vol. 76, no. 4, pp. 740–772, Dec. 2012, doi: 10.1128/MMBR.00035-12.

[66] M. Pohlschroder and R. N. Esquivel, 'Archaeal type IV pili and their involvement in biofilm formation', *Front. Microbiol.*, vol. 6, 2015, Accessed: Feb. 07, 2022. [Online]. Available: https://www.frontiersin.org/article/10.3389/fmicb.2015.00190

[67] K. Lassak, A. Ghosh, and S.-V. Albers, 'Diversity, assembly and regulation of archaeal type IV pili-like and non-type-IV pili-like surface structures', *Res. Microbiol.*, vol. 163, no. 9–10, pp. 630–644, Dec. 2012, doi: 10.1016/j.resmic.2012.10.024.

[68] V. A. Meshcheryakov and M. Wolf, 'Crystal structure of the flagellar accessory protein FlaH of Methanocaldococcus jannaschii suggests a regulatory role in archaeal flagellum assembly', *Protein Sci. Publ. Protein Soc.*, vol. 25, no. 6, pp. 1147–1155, Jun. 2016, doi: 10.1002/pro.2932.

[69] C. J. Schiltz, A. Lee, E. A. Partlow, C. J. Hosford, and J. S. Chappie, 'Structural characterization of Class 2 OLD family nucleases supports a two-metal catalysis mechanism for cleavage', *Nucleic Acids Res.*, vol. 47, no. 17, pp. 9448–9463, Aug. 2019, doi: https://doi.org/10.1093/nar/gkz703.

[70] F. Ito, H. Chishiki, S. Fushinobu, and T. Wakagi, 'Archaeal aldehyde dehydrogenase ST0064 from Sulfolobus tokodaii, a paralog of non-phosphorylating glyceraldehyde-3-phosphate dehydrogenase, is a succinate semialdehyde dehydrogenase', *Biosci. Biotechnol. Biochem.*, vol. 77, no. 6, pp. 1344–1348, 2013, doi: 10.1271/bbb.130119.

[71] F. Blachier, C. Boutry, C. Bos, and D. Tomé, 'Metabolism and functions of L-glutamate in the epithelial cells of the small and large intestines', *Am. J. Clin. Nutr.*, vol. 90, no. 3, pp. 814S-821S, Sep. 2009, doi: 10.3945/ajcn.2009.27462S.

[72] J. J. Van Beeumen, G. Van Driessche, M. Y. Liu, and J. LeGall, 'The primary structure of rubrerythrin, a protein with inorganic pyrophosphatase activity from Desulfovibrio vulgaris. Comparison with hemerythrin and rubredoxin.', *J. Biol. Chem.*, vol. 266, no. 31, pp. 20645–20653, Nov. 1991, doi: 10.1016/S0021-9258(18)54757-8.

[73] I. Lüke *et al.*, 'Biosynthesis of the respiratory formate dehydrogenases from Escherichia coli: characterization of the FdhE protein', *Arch. Microbiol.*, vol. 190, no. 6, pp. 685–696, Dec. 2008, doi: 10.1007/s00203-008-0420-4.

[74] J. P. Cardenas, R. Quatrini, and D. S. Holmes, 'Aerobic Lineage of the Oxidative Stress Response Protein Rubrerythrin Emerged in an Ancient Microaerobic, (Hyper)Thermophilic Environment', *Front. Microbiol.*, vol. 7, 2016, Accessed: Jan. 17, 2022. [Online]. Available: https://www.frontiersin.org/article/10.3389/fmicb.2016.01822

[75] S. Mesnage, 'Bacterial SLH domain proteins are non-covalently anchored to the cell surface via a conserved mechanism involving wall polysaccharide pyruvylation', *EMBO J.*, vol. 19, no. 17, pp. 4473–4484, Sep. 2000, doi: 10.1093/emboj/19.17.4473.

[76] J. D. Esko, T. L. Doering, and C. R. Raetz, 'Eubacteria and Archaea', in *Essentials of Glycobiology*, A. Varki, R. D. Cummings, J. D. Esko, H. H. Freeze, P. Stanley, C. R. Bertozzi, G. W. Hart, and M. E. Etzler, Eds., 2nd ed.Cold Spring Harbor (NY): Cold Spring Harbor Laboratory Press, 2009. Accessed: Jan. 14, 2022. [Online]. Available: http://www.ncbi.nlm.nih.gov/books/NBK1945/

[77] W. A. M. Loenen, D. T. F. Dryden, E. A. Raleigh, G. G. Wilson, and N. E. Murray, 'Highlights of the DNA cutters: a short history of the restriction enzymes', *Nucleic Acids Res.*, vol. 42, no. 1, pp. 3–19, Jan. 2014, doi: 10.1093/nar/gkt990.

[78] N. E. Murray, 'Type I Restriction Systems: Sophisticated Molecular Machines (a Legacy of Bertani and Weigle)', *Microbiol. Mol. Biol. Rev.*, Jun. 2000, doi: 10.1128/MMBR.64.2.412-434.2000.

[79] N. Willemse and C. Schultsz, 'Distribution of Type I Restriction–Modification Systems in Streptococcus suis: An Outlook', *Pathogens*, vol. 5, no. 4, p. 62, Nov. 2016, doi: 10.3390/pathogens5040062.

[80] R. Korona, B. Korona, and B. R. Y. 1993 Levin, 'Sensitivity of naturally occurring coliphages to type I and type II restriction and modification', *Microbiology*, vol. 139, no. 6, pp. 1283–1290, doi: 10.1099/00221287-139-6-1283.

[81] L. Roer, F. M. Aarestrup, and H. Hasman, 'The EcoKI Type I Restriction-Modification System in Escherichia coli Affects but Is Not an Absolute Barrier for Conjugation', *J. Bacteriol.*, vol. 197, no. 2, pp. 337–342, Jan. 2015, doi: 10.1128/JB.02418-14.

[82] M. Chen *et al.*, 'Biochemical and structural characterization of oxygen-sensitive 2-thiouridine synthesis catalyzed by an iron-sulfur protein TtuA', *Proc. Natl. Acad. Sci.*, vol. 114, no. 19, pp. 4954–4959, May 2017, doi: 10.1073/pnas.1615585114.

[83] N. Shigi, 'Biosynthesis and functions of sulfur modifications in tRNA', *Front. Genet.*, vol. 5, 2014, Accessed: Feb. 07, 2022. [Online]. Available: https://www.frontiersin.org/article/10.3389/fgene.2014.00067

[84] C. Wu *et al.*, 'Prokaryotic expression, purification, and production of polyclonal antibody against human polypeptide N-acetylgalactosaminyltransferase 14', *Protein Expr. Purif.*, vol. 56, no. 1, pp. 1–7, Nov. 2007, doi: 10.1016/j.pep.2007.04.027.

[85] M. Dadashipour, M. Iwamoto, M. M. Hossain, J. Akutsu, Z. Zhang, and Y. Kawarabayasi, 'Identification of a Direct Biosynthetic Pathway for UDP–N-Acetylgalactosamine from Glucosamine-6-Phosphate in Thermophilic Crenarchaeon Sulfolobus tokodaii', *J. Bacteriol.*, vol. 200, no. 10, pp. e00048-18, Apr. 2018, doi: 10.1128/JB.00048-18.

[86] R. Tam and J. M H Saier, 'Structural, functional, and evolutionary relationships among extracellular solute-binding receptors of bacteria', *Microbiol. Rev.*, Jun. 1993, doi: 10.1128/mr.57.2.320-346.1993.

[87] W. Saurin and E. Dassa, 'Sequence relationships between integral inner membrane proteins of binding protein-dependent transport systems: Evolution by recurrent gene duplications: Evolution of periplasmic permeases', *Protein Sci.*, vol. 3, no. 2, pp. 325–344, Feb. 1994, doi: 10.1002/pro.5560030216.

[88] R. K. Thauer, A.-K. Kaster, M. Goenrich, M. Schick, T. Hiromoto, and S. Shima, 'Hydrogenases from methanogenic archaea, nickel, a novel cofactor, and H2 storage', *Annu. Rev. Biochem.*, vol. 79, pp. 507–536, 2010, doi: 10.1146/annurev.biochem.030508.152103.

[89] R. D. Mullins, 'Actin-Related Proteins', in *Encyclopedia of Biological Chemistry (Second Edition)*, W. J. Lennarz and M. D. Lane, Eds., Waltham: Academic Press, 2013, pp. 36–41. doi: 10.1016/B978-0-12-378630-2.00468-0.

[90] T. D. Pollard, W. C. Earnshaw, J. Lippincott-Schwartz, and G. T. Johnson, Eds., ‘Chapter 33 - Actin and Actin-Binding Proteins’, in *Cell Biology (Third Edition)*, Elsevier, 2017, pp. 575–591. doi: 10.1016/B978-0-323-34126-4.00033-5.

[91] T. M. Witkos and M. Lowe, ‘The Golgin Family of Coiled-Coil Tethering Proteins’, *Front. Cell Dev. Biol.*, vol. 3, 2016, doi: https://doi.org/10.3389/fcell.2015.00086.

[92] H. Magidovich and J. Eichler, ‘Glycosyltransferases and oligosaccharyltransferases in Archaea: putative components of the N-glycosylation pathway in the third domain of life’, *FEMS Microbiol. Lett.*, vol. 300, no. 1, pp. 122–130, Nov. 2009, doi: 10.1111/j.1574-6968.2009.01775.x.

[93] H. Magidovich and J. Eichler, ‘Glycosyltransferases and oligosaccharyltransferases in Archaea: putative components of the N-glycosylation pathway in the third domain of life’, *FEMS Microbiol. Lett.*, vol. 300, no. 1, pp. 122–130, Nov. 2009, doi: 10.1111/j.1574-6968.2009.01775.x.

[94] A. Anandan and A. Vrielink, ‘Structure and function of lipid A-modifying enzymes’, *Ann. N. Y. Acad. Sci.*, vol. 1459, no. 1, pp. 19–37, Jan. 2020, doi: 10.1111/nyas.14244.

[95] S. D. Breazeale, A. A. Ribeiro, and C. R. H. Raetz, ‘Origin of Lipid A Species Modified with 4-Amino-4-deoxy-l-arabinose in Polymyxin-resistant Mutants of Escherichia coli: AN AMINOTRANSFERASE (ArnB) THAT GENERATES UDP-4-AMINO-4-DEOXY-l-ARABINOSE *’, *J. Biol. Chem.*, vol. 278, no. 27, pp. 24731–24739, Jul. 2003, doi: 10.1074/jbc.M304043200.

[96] A. Mahnert, M. Blohs, M.-R. Pausan, and C. Moissl-Eichinger, ‘The human archaeome: methodological pitfalls and knowledge gaps’, *Emerg. Top. Life Sci.*, vol. 2, no. 4, pp. 469–482, Dec. 2018, doi: 10.1042/ETLS20180037.

[97] A. Dell, A. Galadari, F. Sastre, and P. Hitchen, ‘Similarities and Differences in the Glycosylation Mechanisms in Prokaryotes and Eukaryotes’, *Int. J. Microbiol.*, vol. 2010, p. e148178, Jan. 2011, doi: 10.1155/2010/148178.

[98] S. Voisin *et al.*, ‘Identification and characterization of the unique N-linked glycan common to the flagellins and S-layer glycoprotein of Methanococcus voltae’, *J. Biol. Chem.*, vol. 280, no. 17, pp. 16586–16593, Apr. 2005, doi: 10.1074/jbc.M500329200.

[99] D. J. VanDyke *et al.*, ‘Identification of a putative acetyltransferase gene, MMP0350, which affects proper assembly of both flagella and pili in the archaeon Methanococcus maripaludis’, *J. Bacteriol.*, vol. 190, no. 15, pp. 5300–5307, Aug. 2008, doi: 10.1128/JB.00474-08.

[100] U. Kärcher *et al.*, ‘Primary structure of the heterosaccharide of the surface glycoprotein of Methanothermus fervidus.’, *J. Biol. Chem.*, vol. 268, no. 36, pp. 26821–26826, Dec. 1993, doi: 10.1016/S0021-9258(19)74185-4.

[101] M. Abu-Qarn and J. Eichler, ‘Protein N-glycosylation in Archaea: defining Haloferax volcanii genes involved in S-layer glycoprotein glycosylation’, *Mol. Microbiol.*, vol. 61, no. 2, pp. 511–525, Jul. 2006, doi: 10.1111/j.1365-2958.2006.05252.x.

[102] B. Chaban, S. Voisin, J. Kelly, S. M. Logan, and K. F. Jarrell, ‘Identification of genes involved in the biosynthesis and attachment of Methanococcus voltae N-linked glycans: insight into N-linked glycosylation pathways in Archaea’, *Mol. Microbiol.*, vol. 61, no. 1, pp. 259–268, Jul. 2006, doi: 10.1111/j.1365-2958.2006.05226.x.

[103] U. Zähringer, H. Moll, T. Hettmann, Y. A. Knirel, and G. Schäfer, ‘Cytochrome b558/566 from the archaeon Sulfolobus acidocaldarius has a unique Asn-linked highly branched hexasaccharide chain containing 6-sulfoquinovose’, *Eur. J. Biochem.*, vol. 267, no. 13, pp. 4144–4149, Jul. 2000, doi: 10.1046/j.1432-1327.2000.01446.x.

[104] M. F. Mescher and J. L. Strominger, ‘Purification and characterization of a prokaryotic glycoprotein from the cell envelope of Halobacterium salinarium’, *J. Biol. Chem.*, vol. 251, no. 7, pp. 2005–2014, Apr. 1976.

[105] A. Tamir and J. Eichler, 'N-Glycosylation Is Important for Proper Haloferax volcanii S-Layer Stability and Function', *Appl. Environ. Microbiol.*, vol. 83, no. 6, pp. e03152-16, Mar. 2017, doi: 10.1128/AEM.03152-16.

[106] M. F. Mescher and J. L. Strominger, 'Structural (shape-maintaining) role of the cell surface glycoprotein of Halobacterium salinarium', *Proc. Natl. Acad. Sci. U. S. A.*, vol. 73, no. 8, pp. 2687–2691, Aug. 1976, doi: 10.1073/pnas.73.8.2687.

[107] M. Sumper, E. Berg, R. Mengele, and I. Strobel, 'Primary structure and glycosylation of the S-layer protein of Haloferax volcanii', *J. Bacteriol.*, vol. 172, no. 12, pp. 7111–7118, Dec. 1990, doi: 10.1128/jb.172.12.7111-7118.1990.

[108] L. E. Tailford, E. H. Crost, D. Kavanaugh, and N. Juge, 'Mucin glycan foraging in the human gut microbiome', Mar. 2015, Accessed: Mar. 24, 2022. [Online]. Available: https://www.ncbi.nlm.nih.gov/pmc/articles/PMC4365749/

[109] B. S. Samuel *et al.*, 'Genomic and metabolic adaptations of Methanobrevibacter smithii to the human gut', *Proc. Natl. Acad. Sci. U. S. A.*, vol. 104, no. 25, pp. 10643–10648, Jun. 2007, doi: 10.1073/pnas.0704189104.

[110] H. H. Wandall, M. A. I. Nielsen, S. King-Smith, N. de Haan, and I. Bagdonaite, 'Global functions of O-glycosylation: promises and challenges in O-glycobiology', *FEBS J.*, vol. 288, no. 24, pp. 7183–7212, 2021, doi: 10.1111/febs.16148.

[111] M. Bektas and D. S. Rubenstein, 'The role of intracellular protein O-glycosylation in cell adhesion and disease', *J. Biomed. Res.*, vol. 25, no. 4, pp. 227–236, Jul. 2011, doi: 10.1016/S1674-8301(11)60031-6.

[112] C. R. Torres and G. W. Hart, 'Topography and polypeptide distribution of terminal N-acetylglucosamine residues on the surfaces of intact lymphocytes. Evidence for O-linked GlcNAc', *J. Biol. Chem.*, vol. 259, no. 5, pp. 3308–3317, Mar. 1984.

[113] P. V. den Steen, P. M. Rudd, R. A. Dwek, and G. Opdenakker, 'Concepts and Principles of O-Linked Glycosylation', *Crit. Rev. Biochem. Mol. Biol.*, vol. 33, no. 3, pp. 151–208, Jan. 1998, doi: 10.1080/10409239891204198.

[114] G. D. Havemann, E. M. Sampson, and T. A. Bobik, 'PduA is a shell protein of polyhedral organelles involved in coenzyme B(12)-dependent degradation of 1,2-propanediol in Salmonella enterica serovar typhimurium LT2', *J. Bacteriol.*, vol. 184, no. 5, pp. 1253–1261, Mar. 2002, doi: 10.1128/JB.184.5.1253-1261.2002.

[115] N. W. Kennedy, S. P. Ikonomova, M. Slininger Lee, H. W. Raeder, and D. Tullman-Ercek, 'Self-assembling Shell Proteins PduA and PduJ have Essential and Redundant Roles in Bacterial Microcompartment Assembly', *J. Mol. Biol.*, vol. 433, no. 2, p. 166721, Jan. 2021, doi: 10.1016/j.jmb.2020.11.020.

[116] E. M. Sampson and T. A. Bobik, 'Microcompartments for B12-dependent 1,2-propanediol degradation provide protection from DNA and cellular damage by a reactive metabolic intermediate', *J. Bacteriol.*, vol. 190, no. 8, pp. 2966–2971, Apr. 2008, doi: 10.1128/JB.01925-07.

[117] L.-N. Liu, M. Yang, Y. Sun, and J. Yang, 'Protein stoichiometry, structural plasticity and regulation of bacterial microcompartments', *Curr. Opin. Microbiol.*, vol. 63, pp. 133–141, Oct. 2021, doi: 10.1016/j.mib.2021.07.006.

[118] C. Bräsen, D. Esser, B. Rauch, and B. Siebers, 'Carbohydrate Metabolism in Archaea: Current Insights into Unusual Enzymes and Pathways and Their Regulation', *Microbiol. Mol. Biol. Rev. MMBR*, vol. 78, no. 1, pp. 89–175, Mar. 2014, doi: 10.1128/MMBR.00041-13.

[119] E. Zangelmi, T. Stanković, M. Malatesta, D. Acquotti, K. Pallitsch, and A. Peracchi, 'Discovery of a New, Recurrent Enzyme in Bacterial Phosphonate Degradation: (R)-1-Hydroxy-2-aminoethylphosphonate Ammonia-lyase', *Biochemistry*, vol. 60, no. 15, pp. 1214–1225, Apr. 2021, doi: 10.1021/acs.biochem.1c00092.

[120] W. W. Metcalf *et al.*, ‘Synthesis of Methylphosphonic Acid by Marine Microbes: A Source for Methane in the Aerobic Ocean’, *Science*, vol. 337, no. 6098, pp. 1104–1107, Aug. 2012, doi: 10.1126/science.1219875.

[121] O. A. Sosa, T. J. Burrell, S. T. Wilson, R. K. Foreman, D. M. Karl, and D. J. Repeta, ‘Phosphonate cycling supports methane and ethylene supersaturation in the phosphate-depleted western North Atlantic Ocean’, *Limnol. Oceanogr.*, vol. 65, no. 10, pp. 2443–2459, 2020, doi: 10.1002/lno.11463.

[122] W. W. Metcalf and W. A. van der Donk, ‘Biosynthesis of Phosphonic and Phosphinic Acid Natural Products’, *Annu. Rev. Biochem.*, vol. 78, pp. 65–94, 2009, doi: 10.1146/annurev.biochem.78.091707.100215.

[123] S. Li and G. P. Horsman, ‘An inventory of early branch points in microbial phosphonate biosynthesis’, Biochemistry, preprint, Apr. 2021. doi: 10.1101/2021.04.07.438883.

[124] K.-S. Ju, J. R. Doroghazi, and W. W. Metcalf, ‘Genomics-enabled discovery of phosphonate natural products and their biosynthetic pathways’, *J. Ind. Microbiol. Biotechnol.*, vol. 41, no. 2, pp. 345–356, Feb. 2014, doi: 10.1007/s10295-013-1375-2.

[125] I. Comas, F. González-Candelas, and M. Zúñiga, ‘Unraveling the evolutionary history of the phosphoryl-transfer chain of the phosphoenolpyruvate:phosphotransferase system through phylogenetic analyses and genome context’, *BMC Evol. Biol.*, vol. 8, no. 1, p. 147, May 2008, doi: 10.1186/1471-2148-8-147.

[126] A. C. Keebaugh and J. W. Thomas, ‘The Evolutionary Fate of the Genes Encoding the Purine Catabolic Enzymes in Hominoids, Birds, and Reptiles’, *Mol. Biol. Evol.*, vol. 27, no. 6, pp. 1359–1369, Jun. 2010, doi: 10.1093/molbev/msq022.

[127] L. Carrijo de Oliveira *et al.*, ‘Reenacting the Birth of a Function: Functional Divergence of HIUases and Transthyretins as Inferred by Evolutionary and Biophysical Studies’, *J. Mol. Evol.*, vol. 89, no. 6, pp. 370–383, Jul. 2021, doi: 10.1007/s00239-021-10010-8.

[128] E. DeMoll and L. Tsai, ‘Conversion of purines to xanthine by Methanococcus vannielii’, *Arch. Biochem. Biophys.*, vol. 250, no. 2, pp. 440–445, Nov. 1986, doi: 10.1016/0003-9861(86)90747-2.

[129] Y.-H. Chen, S.-Y. Wang, and J.-C. Hsu, ‘Effect of Caecectomy on Body Weight Gain, Intestinal Characteristics and Enteric Gas Production in Goslings’, *Asian-Australas. J. Anim. Sci.*, vol. 16, no. 7, pp. 1030–1034, Jan. 2003, Accessed: Feb. 15, 2022. [Online]. Available: https://www.animbiosci.org/journal/view.php?doi=10.5713/ajas.2003.1030

[130] R. T. Papke, J. E. Koenig, F. Rodríguez-Valera, and W. F. Doolittle, ‘Frequent recombination in a saltern population of Halorubrum’, *Science*, vol. 306, no. 5703, pp. 1928–1929, Dec. 2004, doi: 10.1126/science.1103289.

[131] H. Cadillo-Quiroz *et al.*, ‘Patterns of gene flow define species of thermophilic Archaea’, *PLoS Biol.*, vol. 10, no. 2, p. e1001265, Feb. 2012, doi: 10.1371/journal.pbio.1001265.

[132] A. Wagner *et al.*, ‘Mechanisms of gene flow in archaea’, *Nat. Rev. Microbiol.*, vol. 15, no. 8, pp. 492–501, Aug. 2017, doi: 10.1038/nrmicro.2017.41.

[133] P. Deschamps, Y. Zivanovic, D. Moreira, F. Rodriguez-Valera, and P. López-García, ‘Pangenome Evidence for Extensive Interdomain Horizontal Transfer Affecting Lineage Core and Shell Genes in Uncultured Planktonic Thaumarchaeota and Euryarchaeota’, *Genome Biol. Evol.*, vol. 6, no. 7, pp. 1549–1563, Jul. 2014, doi: 10.1093/gbe/evu127.

[134] E. E. Hansen *et al.*, ‘Pan-genome of the dominant human gut-associated archaeon, Methanobrevibacter smithii, studied in twins’, *Proc. Natl. Acad. Sci. U. S. A.*, vol. 108 Suppl 1, pp. 4599–4606, Mar. 2011, doi: 10.1073/pnas.1000071108.
